# An ancient gene regulatory network sets the position of the forebrain in chordates

**DOI:** 10.1101/2023.03.13.532359

**Authors:** Giacomo Gattoni, Daniel Keitley, Ashley Sawle, Elia Benito-Gutiérrez

## Abstract

The evolutionary origin of the vertebrate brain is still a major subject of debate. Its distinctive dorsal position and development from a tubular neuroepithelium are unique to the chordate phylum. Conversely, apical organs (AO) are larval sensory/neurosecretory centers found in many invertebrate taxa, including in animals without a brain. Previous studies have shown that AOs are specified by a conserved set of genes under the influence of Wnt signalling. Although most of these genes are expressed in chordate nervous systems (including vertebrates), no AOs have ever been described in this group of animals. Here we have exploited single-cell genomic approaches to characterize cells showing AO profiles in sea urchin (ambulacrarian), amphioxus (invertebrate chordate) and zebrafish (vertebrate chordate). This, in combination with co-expression analysis in amphioxus embryos, has allowed us to identify an active and dynamic anterior Gene Regulatory Network (aGRN) in the three deuterostome species. We have further discovered that as well as controlling AO specification in sea urchin, this aGRN is involved in the formation of the hypothalamic region in amphioxus and zebrafish. Using a functional approach, we find that the aGRN is controlled by Wnt signalling in amphioxus, and that suppression of the aGRN by Wnt overactivation leads to a loss of forebrain cell types. The loss of the forebrain does not equate to a reduction of neuronal tissue, but to a loss of identity, suggesting a new role for Wnt in amphioxus in specifically positioning the forebrain. We thus propose that the aGRN is conserved throughout bilaterians and that in the chordate lineage was incorporated into the process of neurulation to position the brain, thereby linking the evolution of the AO to that of the chordate forebrain.

## Introduction

The vertebrate brain is one of the most complex structures in nature. Despite significant advancements in our understanding of brain anatomy and physiology, its evolutionary origin remains a major subject of debate. While complex brains have been described in protostome (anellids, arthropods, mollusks) and deuterostome (chordates) taxa, they display significant differences in architecture and development^1^. Moreover, establishing a link between the central nervous system (CNS) of these distantly related groups has been particularly challenging as chordates are the only deuterostome group possessing a brain^2,3^. The other two deuterostome phyla, echinoderms and hemichordates, lack an identifiable brain, and the derived pentaradial organization of the echinoderm body makes evolutionary comparisons challenging^4,5^. However, a common aspect of nervous system development in ambulacrarians (echinoderms and hemichordates), also shared with protostomes^6–12^ and cnidarians^13–15^, is the presence of larval apical organs (AO)^16–18^. These are sensory-neurosecretory structures that develop from the anteriormost part of the ectoderm in all these phyla^19,20^, suggesting that the apical gene regulatory network (aGRN) specifying AOs might have appeared with the ancestor of Eumetazoa (Cnidaria + Bilateria (Protostomes + Deuterostomes))^20–22^.

AOs were initially defined by the presence of serotonergic neurons^9,18,23^ but, more recently, a shared set of transcription factors expressed in AO cells has been identified^11,21,24–26^. The common denominator in AO specification in cnidarians, protostomes and deuterostomes is the early mutually repressive interaction between Wnt/β-catenin signalling emanating from the future blastoporal side of the embryo (vegetal pole for Bilateria, oral pole in Cnidaria) and the transcription factors *Six3/6* and *FoxQ2*, localized on the opposite side (animal pole in Bilateria, aboral pole in Cnidaria)^11,27–31^. These repressive interactions have been demonstrated by knock-down experiments and pharmacological treatments in cnidarian, protostome and ambulacrarian larvae. In all cases, Wnt overactivation results in a loss of *Six3/6* and *FoxQ2* expression from the aboral/animal pole, followed by the loss of the AO^11,31–34^. Conversely, Wnt inhibition leads to an expansion of the *Six3/6* and *FoxQ2* domain across ectoderm, which consequently is entirely specified as anterior neuroectoderm (ANE)^34–40^. This data supports a model of apical/anterior patterning in which members of the Wnt family repress the expression of *Six3/6* and *FoxQ2* at the vegetal/oral pole, while *Six3/6* represses Wnt expression at the animal/aboral pole and acts together with *FoxQ2* to specify the apical/aboral region^41,42^. Later in the development of ambulacrarians, after the establishment of the animal (future anterior) and vegetal (future posterior) domains during cleavage, non-canonical Wnt pathways progressively restrict *Six3/6* and *FoxQ2* anteriorly towards the animal pole via interactions with the Wnt receptor *Frz5/*8^31,36^. At the animal pole, Wnt inhibitors of the sFRP and Dkk families prevent *Six3/6* and *FoxQ2* downregulation by Wnt^31,34,43,44^. These interactions result in the correct positioning and temporal synchronization of downstream transcription factors that are crucial for AO development in many bilaterian invertebrate larvae^21,45^.

AOs have never been described in chordates. Nonetheless, the genes involved in AO development are also present in chordate genomes and many of them operate during the specification of the anterior nervous system^46^. *Six3*, for example, is one of the earliest genes expressed in the anterior neural plate of vertebrate embryos^47,48^. Six3 is essential for the formation of the vertebrate secondary prosencephalon (telencephalon and hypothalamus) and is required to inhibit the specification of diencephalic cell types anteriorly^49,50^. The mutually repressive interactions between Six3 and Wnt have been also demonstrated in vertebrate nervous systems, where they are essential to establish the anterior identity of the brain^49^. However, unlike in the AO of invertebrate larvae, *Six3* expression in vertebrates is not influenced by *FoxQ2*, which has been lost in placental mammals^51,52^. Adding to this striking difference, *Six3* in vertebrates is expressed well after gastrulation has initiated in cells already committed to a neuronal fate^53,54^. While it is assumed that the specification of the ANE/ANB (Anterior neural border) implies that anterior ectodermal cells acquire a neural identity, little is known about the mechanism that transforms anterior ectodermal cells into neurectoderm, and if Six3 is involved in this process.^55^.

Considering that several components of the GRN that specifies the AO are involved in the development of the vertebrate brain, it has been proposed that the anterior brain evolved in bilaterians by coalescence of two separated neuronal systems: an anterior ‘apical nervous system’ (ANS) and a posterior ‘blastoporal nervous system’ (BNS). This chimeric brain hypothesis^56^, very similar to the earlier “animal-axial” hypothesis^57^, proposes that the ANS corresponds to the AO of invertebrate larvae, while the BNS corresponds to the remaining nervous system (posterior to the AO). However, both hypotheses pose the same fundamental problem regarding the conflicting fate of anterior neuroectodermal precursors in ambulacrarians and chordates: in ambulacrarians the ANE forms the AO^58^, while in chordates the ANE gives rise to the forebrain^21,47,59^.

To shed light into the evolutionary link between invertebrate AOs and chordate brains, we focused on the invertebrate chordate amphioxus, which branches at the base of the chordate phylogenetic tree and for which there is evidence of an anterior domain expressing *Six3/6* and *FoxQ2* early in gastrulation^60–62^. We first surveyed publicly available scRNA-seq datasets from sea urchin embryos^63^ to corroborate genes distinctively involved in AO formation and defined those in common with the vertebrate forebrain. We then co-profiled in amphioxus, by RNA-seq and *in situ* hybridization, the expression of *Six3/6, FoxQ2* and Wnt inhibitors, as well as several downstream genes that we identify here as a part of a sequential cascade characterizing early and late phases of AO development in sea urchin. Our results demonstrate that the apical (anterior) region of early amphioxus embryos is similarly specified to that of the sea urchin AO. Using a functional approach, we find that this apical region in amphioxus is also controlled by mutually repressive interactions between the Wnt/β-catenin signalling pathway and the transcription factors *Six3/6* and *FoxQ2*. This indicates that the aGRN is conserved in amphioxus despite lacking an AO. Later in development, during neurulation, the expression of most of these genes restricts, with exquisite precision, to specify anterior parts of the amphioxus brain. We identify this region as homologous to the vertebrate retina and hypothalamus based on shared gene expression profiles. Furthermore, our results show that failure to specify the anterior region by ectopic activation of the Wnt/β-catenin signalling pathway in amphioxus results in a loss of retinal- and hypothalamic-like precursors. Altogether, our work shows that the ANE in bilaterians is specified by a conserved aGRN, which controls the differentiation of the AO in ambulacrarian larvae and the retino-hypothalamic region of the amphioxus brain, thereby revealing the evolutionary link between the invertebrate AO and the chordate forebrain.

## Results

### The AO in sea urchins is specified through bi-phasic co-expression of a particular set of genes that in zebrafish operate collectively during the development of the hypothalamus

The animal pole of many invertebrate embryos is set by an anterior (or apical) GRN (aGRN) directing the formation of the ANE and the subsequent specification of larval AOs ^21,22^. Although AOs have never been described in vertebrates, a possible homology between ambulacrarian and vertebrate ANEs has been previously proposed based on molecular evidence^21^. However, the ANE precursors in these two lineages have distinct fates, giving rise to the AO in ambulacrarians and the forebrain in vertebrates. To investigate whether a similar profile is recognizable in the ANE derivatives of both lineages, we first sought to define the co-expression dynamics of genes involved in the development of the ambulacrarian AO and the vertebrate forebrain, at a single-cell level. To this end, we comparatively analyzed publicly available scRNAseq datasets from the sea urchin (*Strongylocentrotus purpuratus*, Echinodermata)^63^ and from zebrafish (*Danio rerio*, Chordata)^64^.

In the sea urchin, the expression and function of apical patterning genes has been well characterized^21,31,37,43,65,66^. The aGRN is composed of at least 11 genes expressed on the animal half of the embryo: *FoxQ2, Six3/6, Frz5/8, sFRP1/2/5a, Otx, Dkk3, Dkk1, Fezf, Rx, Nk2*.*1* and *Lhx2/9*^11,21^. We used this set of genes to query the scRNA-seq dataset published by Foster and colleagues. The original analysis identified two neuronal clusters enriched for FoxQ2 expression, but these were not described in relation to the other genes in the aGRN, nor as belonging to the AO. Therefore, we performed a finer re-annotation of AO cell types in the Foster et al. dataset, by querying the expression of aGRN genes and the serotonergic neurons marker, *Tph*, across developmental stages (see Materials and Methods). We find a clear temporal sequence of gene activation, divisible into early and late phases (Figure 1A and Supplementary Figure 1A-B). In particular, *Six3/6* and *FoxQ2* start to be expressed zygotically at the morula stage, together with the maternally expressed *Otx, Frz5/8* and *sFRP1/2/5* (Supplementary Figure 1C*). Dkk3* starts to be transcribed at the morula stage followed by *Dkk1*, and both are strongly expressed by the hatched blastula stage. *Fezf, Rx, Nk2*.*1* and *Lhx2/9* are activated next and gradually increase their expression throughout the mesenchyme blastula and early gastrula stages. By the end of gastrulation, we see the first differentiating serotonergic cells starting to express *Tph* (Supplementary Figure 1C).

**Figure 1.**
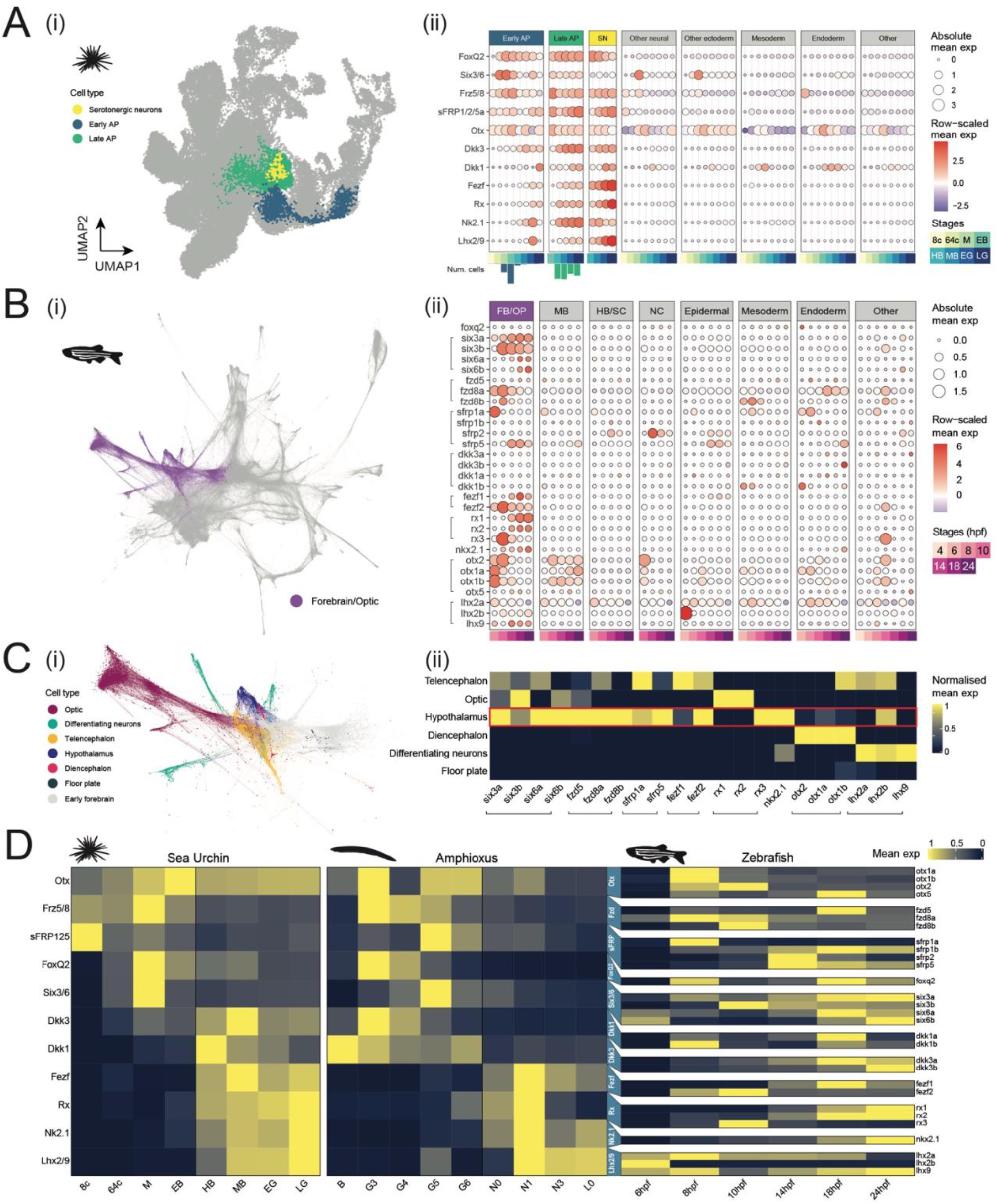
Anterior gene regulatory network (aGRN) signature in deuterostome scRNAseq data. **A** i) Uniform manifold approximation and projection (UMAP) plot of sea urchin single-cell transcriptomics data reanalyzed from Foster et al. 2020, highlighting apical plate (AP) cell types, giving rise to the AO. ii) Dotplot showing the mean expression of aGRN genes across sea urchin cell types through developmental time. Normalized (z-score) values across each gene are represented by the dot color, allowing for comparisons between genes with different absolute mean levels of expression (represented by dot size). The bar plots annotating early and late apical plate cell types show the relative proportion of cells across developmental stages and confirm that early and late apical patterning is mostly restricted to early and late timepoints respectively. **B** i) Force-directed graph layout of zebrafish single-cell data from Wagner et al. 2018, highlighting forebrain/optic cell types as a match to the sea urchin AP cell types. ii) Mean expression of apical patterning gene homologs across zebrafish cell types through developmental time highlight a broad apical patterning signature in the zebrafish forebrain. Brackets highlight paralog groups. **C** i) Zebrafish forebrain/optic cells of the force-directed graph layout in Bi, colored according to a deeper reannotation of cell types. A small number of outlier cells are not shown for aesthetic purposes. ii) aGRN homologs expressed across refined forebrain/optic cell types show that the strongest signature is observed in the zebrafish hypothalamus (highlighted in red). Min-max normalization is performed across genes within each cell type. 8c: 8-cell stage; 64c:64 cell stage; M: morula; EB: early blastula; MB: mesenchymal blastula; EG: Early gastrula; LG: late gastrula. Zebrafish stages given in terms of hours post-fertilization. **D** Comparison of aGRN activation in sea urchin, zebrafish and amphioxus. Normalized mean expression of apical patterning genes within neural tissues across developmental stages in the sea urchin (based on Foster et al. 2020), amphioxus (based on Ma et al. 2022) and zebrafish (based on Wagner et al 2018). Black vertical lines separate an early and late phase of apical patterning in the sea urchin and amphioxus. Blue trapeziums connect the one-to-many homologs between sea urchin/amphioxus and zebrafish apical patterning genes. B: blastula; G3: early gastrula - G6: late gastrula; N1: early neurula – N3: mid neurula; L0: larval stage.

Although some of these genes are independently expressed across many different tissues (Supplementary Figure 1D), their combination is spatially restricted to this group of cells that we annotate as the AO (Figure 1A). Within this AO mega cluster, cells follow a trajectory of maturation that recapitulates the sequential gene activation observed above. Accordingly, these cells can be divided into three groups representing an early and a late phase of specification, and a phase of terminal differentiation. The cells that we define as having an “early apical plate” (EAP) identity predominantly belong to embryos at developmental stages between the morula and the early blastula stage, and they are characterized by the expression of *FoxQ2, Six3/6, Frz5/8, sFRP1/2/5a, Dkk3* and *Otx*. The cells that we define as having a “late apical plate” (LAP) identity belong mostly to embryos at developmental stages between the hatched blastula and the late gastrula stages and add the expression of *Dkk1, Fezf, Otx, Rx, Nk2*.*1*, and *Lhx2/9* to the EAP profile (Figure 1Ai, Aii, and Supplementary Figure 1B and C).

Within the LAP cluster, we identify the serotonergic neurons of the AO as a group of cells that only at the late gastrula stage start expressing *Tph* (Supplementary Figure 1C and D). This cluster of serotonergic neurons is also marked by strong *Fezf* and *Lhx2/9* expression (Figure 1Aii), but have *Six3/6* switched off, and low levels of most early genes. Overall, our analysis in the sea urchin reveals a clear sequence in the activation of apical patterning genes that ties with the progress of cell differentiation and neurogenesis in the apical organ.

Using the same set of 11 genes, we queried a time-course scRNA-seq dataset generated from zebrafish embryos across 4-24 hours post-fertilization (hpf)^67^. At least one paralog for each orthologous gene, apart from FoxQ2, is spatially restricted to a particular region of the force-directed graph embedding. In the original publication, these cells are annotated as a broad brain region encompassing the forebrain and optic cups (Figure 1Bi). Although not all paralogs for each gene are expressed there, at least one of the orthologs of *Six3/6, Frz5/8, sFRP1/2/5, Fezf, Rx, Nkx2*.*1, Otx* and *Lhx2/9* are localized to this cluster (Figure 1Bii).

To gain a better insight on the localization of these cells within the forebrain cluster, we used definitions given by the prosomeric model, which considers the hypothalamus as part of the “secondary prosencephalon” (the most anterior/rostral part of the brain) and separated from the diencephalon^68,69^. In the original study, Wagner et al. annotate telencephalon and diencephalon subclusters within the forebrain/optic region; however, no distinction was made between the hypothalamus and diencephalon. We therefore isolated the cells originally annotated as “diencephalon” and queried them for transcription factors that are known to regionalize this brain region at these developmental stages^53,70–72^. This allowed us to distinguish hypothalamic cell types, which express *Fezf1, Fezf2, Nk2*.*4* and *Rx3*, from diencephalic cell types expressing *Barhl2, Irx3a, Pitx2* and *Pitx3* (Supplementary Figure 1E-F). We then used the re-annotated dataset to plot our set of apical patterning transcription factors across specific forebrain cell types (Figure 1C). This revealed that an apical patterning signature is present within the hypothalamus, and, to a lesser extent, in the retina and the telencephalon, while it is absent from the posterior diencephalon and the floor plate (Figure 1Cii). Altogether, our results identify a conserved set of 10 genes that act collectively during the formation of the apical organ in sea urchins and during the specification of the hypothalamic region in zebrafish.

### The aGRN in amphioxus involves the expression of the conserved set of genes specifying the sea urchin AO and the zebrafish hypothalamus

Having identified a shared set of genes involved in the specification of the ANE in sea urchin and zebrafish, we set out to characterize the expression dynamics of each of the orthologous genes during the development of amphioxus. We first analyzed a recently published single-nuclei RNAseq dataset of *B. floridae* development^73^ to compare the temporal activation of each ortholog in sea urchin, amphioxus and zebrafish (Figure 1D and Supplementary Figure 2). By looking at the dynamics of expression in ectodermal tissues of each species, we detect a clear biphasic separation in amphioxus, which is similar to sea urchin: EAP genes such as *FoxQ2, Six3/6, Frz5/8, sFRP1/2/5* were expressed early in amphioxus, while late genes *Fezf, Rx, Nk2*.*1* and *Lhx2/9* started to be expressed only at the beginning of neurulation. Compared to sea urchin, however, the two *Dkk* orthologs were expressed at earlier stages. Conversely, no clear separation between early and late phases was observed in zebrafish (Figure 1D).

Similar results were obtained by analyzing a published RNAseq dataset of *B. lanceolatum* spanning from cleavage to larval stages (Supplementary Figure 3)^74^. Differential expression analyses of all aGRN genes at these different developmental time points revealed that, as in the sea urchin, the first transcription factor to be activated is *FoxQ2* (Supplementary Figure 3). We find *FoxQ2* simultaneously expressed with *Wnt8* at the blastula stage. The expression of both genes peaks during gastrulation and then slowly decreases over developmental time, still maintaining high levels of expression during gastrulation and up to mid neurulation. During gastrulation, *Six3/*6, *Otx, Frz5/8* and *sFRP1/2/5a* become strongly expressed, (Supplementary Figure 2B), closely resembling the early phase of specification observed in the sea urchin. During early neurula stages, *Fezf, Nk2*.*1* and then *Rx* and *Lhx2/9b* are upregulated as in the late phase of AO specification in sea urchin. We see the expression of most “early” genes decreasing at later larval stages, implying a tight temporal regulation (Supplementary Figure 3). Overall, we find that amphioxus has an active aGRN with biphasic dynamics similar to that observed in sea urchin embryos.

We then set out to spatially resolve the expression and co-expression of all genes described above, at the developmental stages when we find them expressed in bulk. We did this by *in situ* hybridization, using our recently optimized protocol based on the chain reaction technique (HCR), which allows the simultaneous detection of multiple genes per embryo^75,76^. We first applied it to visualize *in situ* the expression of *FoxQ2* and *Wnt8* from the blastula to the mid-neurula stage when significant levels of both genes were detected in our RNA-seq results (Supplementary Figure 3). These two genes were co-profiled together with the broad early neural marker Neurogenin (*Ngn*), to control for neuronal tissue specification (Figure 2A). *FoxQ2* was first detected at the 128-cell stage in the nuclei of cells in the animal half of the embryo and became more defined at the B (blastula) stage (Figure 2A, Supplementary Figure 4). These *FoxQ2* positive cells were bordered at the equator of the embryos by weak nuclear *Wnt8* expression. We observed this complementary pattern persisting until early gastrula stages (G3), when *Wnt8* labels the blastoporal lips (Figure 2A). At this stage, *Ngn* starts to be visible in a dorsal band of cells within the *FoxQ2* domain abutting the Wnt8-positive cells at the dorsal blastoporal lip (Figure 2A). At the G4 stage (late gastrula, 8hpf), the expression of *FoxQ2* starts to restrict towards the anterior side of the embryo and over time it becomes slightly dorsalized and absent from the ventral side. At the same time, *Wnt8* restricts to the lateral and ventral mesendoderm. The dorsal *Ngn*-positive band instead expands at the expense of *FoxQ2*, the expression of which restricts antero-dorsally (Figure 2A, Supplementary Figure 4). Expression of *FoxQ2* continues to restrict through neurulation, such that by the 7-somites stage (7ss) it only occupies the anterior most tip of the ectoderm, where it no longer co-localizes with *Ngn*, rendering it absent from the neuroectoderm (Figure 2A, Supplementary Figure 4). Both the relative volume and relative number of *FoxQ2*+ cells significantly decrease during this time (Supplementary Figure 4), demonstrating that the initial expression of *FoxQ2* across the animal half of the blastula and its restriction during gastrulation resembles the establishment of animal and vegetal fields and the early apical patterning of ambulacrarian larvae.

**Figure 2.**
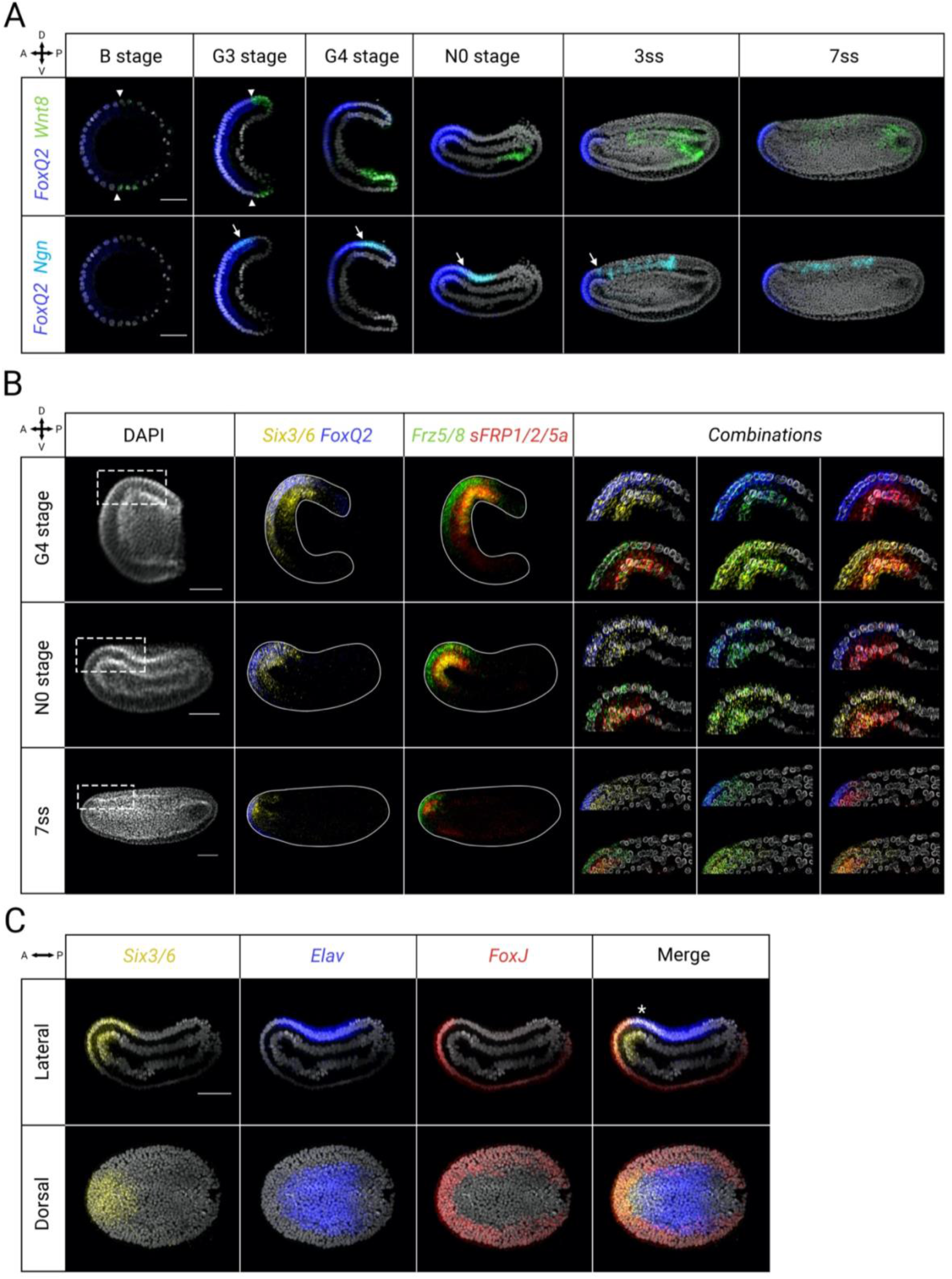
Early aGRN in amphioxus embryogenesis visualized through *in situ* HCR. **A** Co-detection of *FoxQ2* (blue) with *Wnt8* (green) and early neural marker *Ngn* (cyan) from blastula to the 7-somites (7ss) stage. Early *FoxQ2* is expressed broadly through the animal side of the embryo in a complementary manner with Wnt8 (arrowheads show expression boundary), but it restricts anteriorly and dorsally during neurulation. The early neural plate (marked by *Ngn*) is initially located within the *FoxQ2* expression domain (arrows point to area of co-expression) until FoxQ2 is downregulated in the nervous system at the 7ss stage. **B** Co-expression and anterior restriction of early aGRN genes during neurulation. Magnified view of the dashed boxes in the DAPI channel as a merge for all genes. *FoxQ2* (blue) is co-expressed with *Six3/6* (yellow) and *Frz5/8* (green) in the late gastrula (G4), while *sFRP1/2/5a* (red) is found dorsally within the domain of the other three genes. During neurulation *FoxQ2, Six3/6* and *Frz5/8* mark progressively smaller areas in the anterior end of the embryo. *sFRP1/2/5a* remains expressed dorsally at the boundary of neural and non-neural ectoderm. **C** Early aGRN gene *Six3/6* (yellow) is expressed in both neural and non-neural ectoderm, as indicated by co-detection with pan-neural marker *Elav* (blue) and ciliated cell marker *FoxJ* (red) in early neurulas (N0). Asterisk indicate neural expression of *Six3/6*. All scale bars are 50µm.

We next searched for EAP identities in amphioxus embryos by co-profiling the expression of four genes acting in the early phase of specification of the AO in the sea urchin (*FoxQ2, Six3/6, Frz5/8* and *sFRP1/2/5a*) (Figure 1, Figure 2B). All genes were expressed in the anterior-dorsal ectoderm at G4 stage (Figure 2B). While *FoxQ2, Six3/6* and *Frz5/8* were co-expressed in a broad antero-dorsal ectodermal area, *sFRP1/2/5a* appeared dorsally restricted within this co-expression domain. At the early and mid-neurula stages *Six3/6* and *Frz5/8* gradually restrict their domains of expression following a similar trend to that observed for *FoxQ2*. Unlike *FoxQ2* however, *Six3/6* and *Frz5/8* remain in the nervous system. At 7ss, *Frz5/8* marks the anterior most part of the neural tube, while *Six3/6* expression is separated into an anterior and posterior population of cells separated by an intercalated *Six3/6*-negative region, as previously described^77^. During the phase of *Six3/6* and *Frz5/8* restriction, *sFRP1/2/5a* remains expressed at the boundary between the anterior neural tube and non-neural ectoderm (Figure 2B). At the same stages considered above, *Dkk3* is also expressed in the anterior ectoderm, similar to *sFRP1/2/5a* (Supplementary Figure 5). This reveals a timing and position tentatively congruent with a role of *sFRP1/2/5a* and *Dkk3* in blocking Wnt signalling anteriorly, as described in the sea urchin^31,58^. Moreover, at the gastrula stage *Otx* is expressed in the antero-dorsal portion of the embryo, corresponding to the anterior portion of the neural plate and including the entire neural expression of *Six3/6* (Supplementary Figure 5). The pattern remained similar through neurulation, when *Otx* embraced both anterior and posterior *Six3/6* domains, reaching out beyond the latter, and filled the intercalated *Six3/6*-negative gap (Supplementary Figure 5).

A distinctive feature of AO patterning in echinoderms is that the initial phase of specification, defined above by the expression of genes such as *Six3/6, FoxQ2* and *Frz5/8*, occurs on progenitor cells that will give rise to both neural and non-neural ectoderm. It is only during the late phase of apical patterning that a subset of these ectodermal progenitors become specified as AO neurons. This contrasts with vertebrates, where we first detect the apical patterning signature within the forebrain, when the distinction between non-neuronal and neuronal ectoderm has been already made (Figure 1). To understand how this is reflected in amphioxus, we co-profiled *Six3/6* with the pan-neural marker *Elav* and the ciliated ectodermal cell marker *FoxJ* (Figure 2C). In neurulating amphioxus embryos *Elav* and *FoxJ* are mutually exclusive, thereby providing us with an opportunity to clearly discern neuronal and non-neural ectoderm (Figure 2C). At these stages, we find *Six3/6* expressed both in anterior-dorsal *Elav*-positive cells and in the anteriorly abutting *FoxJ*-positive cells. As development progresses and the neural tube becomes morphologically apparent, *Six3/6* restricts anteriorly but continues to be expressed in both neural and non-neural anterior ectoderm (Supplementary Figure 5). Altogether, our results indicate that apical patterning in amphioxus has an initial phase of specification acting on anterior ectodermal cells, including those fated to neuronal and non-neural identities, and before any neuronal differentiation is observed. This strongly resembles the condition in ambulacrarians, suggesting that this initial phase of apical patterning is conserved in basally branching chordates.

### Late AP genes are expressed in the amphioxus brain

We next focused on spatially resolving late AP (LAP) identities in amphioxus embryos, by co-profiling the expression of genes active in the late phase of AO specification in sea urchin (*Dkk1, Fezf, Otx, Rx, Nk2*.*1*, and *Lhx2/9)* together with *Six3/6*, which is active throughout the two phases (Figure 1D, Figure 3). Given the strong aGRN signal obtained from the zebrafish forebrain, we included in this analysis amphioxus embryos at later developmental stages, when the cerebral vesicle is morphologically visible (Figure 3).

**Figure 3.**
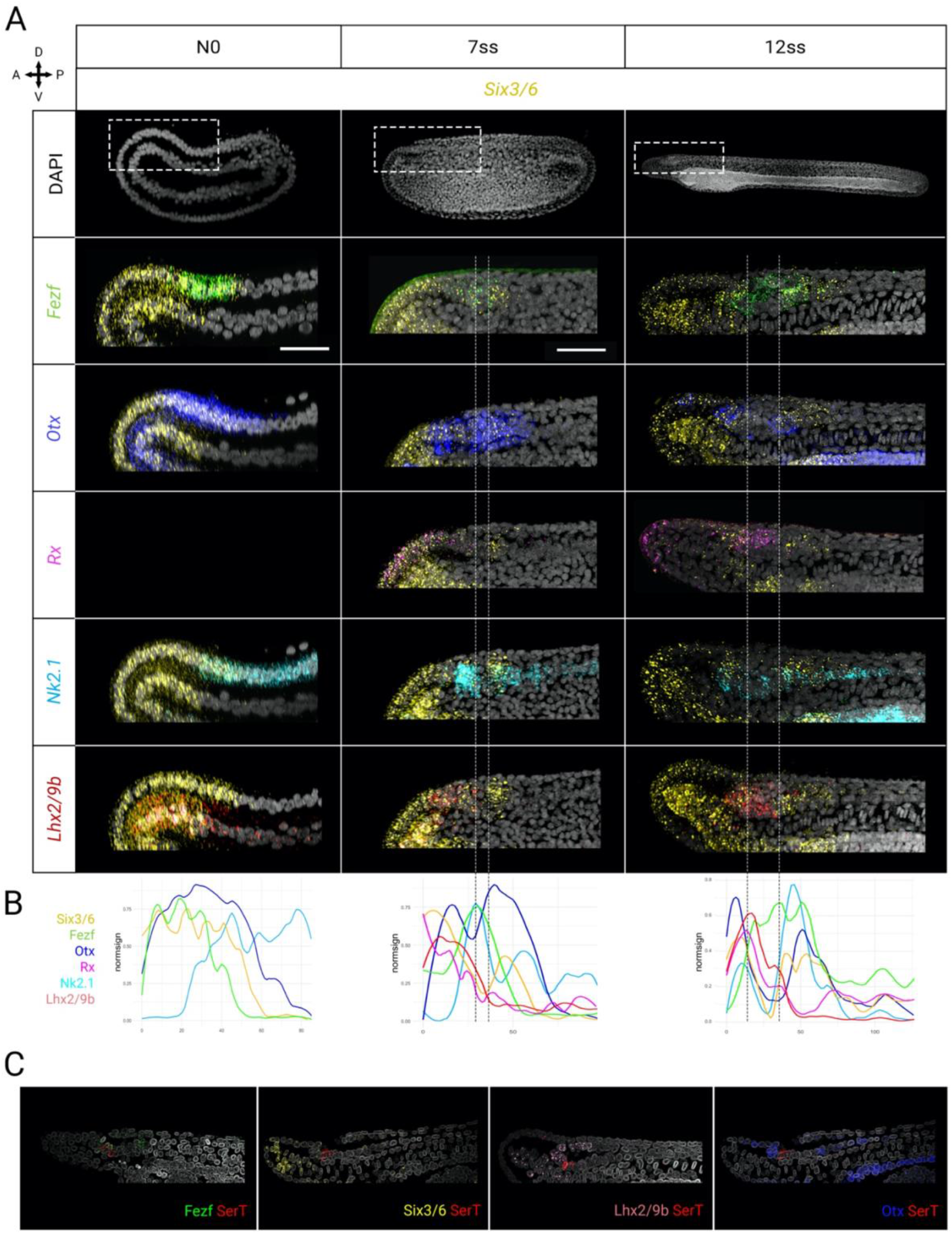
Late aGRN genes are expressed in the anterior brain during amphioxus neurulation. **A** *Six3/6* (yellow) co-expression with core set of late aGRN genes through neurulation, from early neurula (N0) to early larva (12ss). Apical organ transcription factors progressively appear in the anterior nervous system. *Fezf* (green) and *Otx* (blue) are broadly expressed in the *Six3/6*-positive domain at N0, *Rx* (magenta) is not active, *Nk2*.*1* (cyan) marks the floor plate and *Lhx2/9b* (red) is expressed only in the mesoderm. At 7ss *Six3/6* expression in the brain splits into two domains separated by an intercalated *Six3/6*-negative region. Rx appears within the *Six3/6* expression domain in both neural and non-neural ectoderm. *Nk2*.*1* and *Lhx2/9* are also expressed in the anterior nervous system anterior to the posterior *Six3/6*-positive region. At 12ss the intercalated *Six3/6*-negative region expands and late aGRN genes remain expressed within the anterior cerebral vesicle. **B** Expression plots showing relative position of each gene along the AP axis within the amphioxus neuroectoderm at the different developmental stages in A. **C** Profiling of serotonergic neurons in the amphioxus early larva (14ss). Serotonergic neurons in the cerebral vesicle, marked by *SerT* (red), are located at the boundary between the anterior *Six3/6*-positive and *Six3/6*-negative region, posterior to *Otx* (blue). They co-express *Fezf* (green) and *Lhx2/9* (pink). Scalebars are 50µm.

In general, LAP markers progressively appear in the anterior-dorsal ectoderm during neurulation, so that by the 12ss stage they are all expressed in the amphioxus cerebral vesicle (Figure 3A-B). *Dkk1* is already visible at G4, but at this stage we find it localized to the anterior mesoderm. Later, at the 7ss, *Dkk1* is detected in the ectoderm, at the border of the future cerebral vesicle (Supplementary Figure 5(4)). *Fezf* starts to be expressed at the beginning of neurulation, when we see it localized in the anterior tip of the neural plate. Subsequently, by the end of neurulation (12ss in Figure 3A-B), *Fezf* expression becomes restricted to individual cells posterior to the frontal eye, the dorsal roof of the cerebral vesicle and posterior to the infundibular organ, where it is co-expressed with *Six3/6* in its posterior domain. At this stage, *Otx* also becomes restricted to individual cells in the brain, including the frontal eye, the roof of the cerebral vesicle and posterior to the infundibular cells, beyond the posterior Six3/6 domain. At the 7ss stage the first sign of *Rx* expression is seen in the anterior ectoderm and at the rostral end of the neural tube, where it continues to be expressed after the formation of the cerebral vesicle. *Nk2*.*1* is seen in floor plate progenitors from the late gastrula stage, but by 7ss, a new population appears in the anterior half of the brain, composed of cells that disperse as the brain elongates through development, as shown at 12ss (Figure 3A-B). By the end of neurulation also a few cells of the frontal eye complex start to express *Nk2*.*1*. Finally, whilst the earliest expression of *Lhx2/9b* is seen in the anterior endoderm, we see it expressed in the brain in embryos at the 7ss, a pattern that is maintained thereafter and through development^78^.

In summary, our results show that genes acting in the late phase of AO specification in echinoderms are predominantly expressed in the anterior ectoderm and the early developing brain in amphioxus.

A well-known feature of AOs in marine invertebrate larvae is the presence of serotonergic neurons. It has been long known that the frontal eye complex in amphioxus is equipped with serotonergic neurons^79,80^, but little is known about the profile of these cells. Having found that in the sea urchin AO serotonergic cells commit to this fate at the end of the late phase of specification and strongly co-express *Fezf* and *Lhx2/9*, we set out to probe amphioxus embryos with the same set of genes. Accordingly, we co-profiled *SerT* in amphioxus at the 14ss stage together with two aGRN genes that continue to be expressed late, *Six3/6* and *Otx*, as well as with two late genes, *Fezf* and *Lhx2/9* (Figure 3C). As in sea urchin, we found that the serotonergic neurons in the amphioxus frontal eye complex are characterized by strong *Fezf* and weak *Lhx2/9* expression but are devoid of *Otx* as previously suggested^81^ and rarely contain *Six3/6* transcripts in their soma (Figure 2C). Altogether, we see that in amphioxus LAP genes are activated in similar domains and follow a similar temporal sequence as in the sea urchin, but in addition, they also contribute to define the increasingly diverse cell type repertoire of the brain.

### The amphioxus aGRN is dependent on Wnt signalling

Several studies have demonstrated that the aGRN specifying the AOs of invertebrate larvae and the ANE of vertebrate embryos is regulated by the Wnt/β-catenin signalling^11,21,47^. Whether this means there is a common underlying mechanism is still an open question. In cnidarian, protostome and deuterostome larvae the expression of *Six3/6* and *FoxQ2* is negatively regulated by members of the Wnt family, in particular Wnt1 and Wnt8, such that ectopic activation of Wnt/β-catenin signalling results in the loss of aGRN genes^11,31,32,34,35^. In vertebrates, Six3 is essential to keep the prospective forebrain region free from inhibitory Wnt signalling, and ectopic activation of Wnt/β-catenin signalling results in a loss of forebrain structures^49^. Given this, we sought to investigate whether the aGRN in amphioxus is also under the control of Wnt/β-catenin signalling. To this aim, we treated embryos with Azakenpaullone (AZA), a strong GSK-3 inhibitor commonly used to over activate Wnt/β-catenin signalling^82^. Developing embryos were treated with either 10µM of AZA or DMSO as a control at two developmental timepoints: from the blastula (B, 4.5hpf) to the early neurula stage (N0, 12hpf); and from the late gastrula until the 7ss stage (Figure 4A).

**Figure 4.**
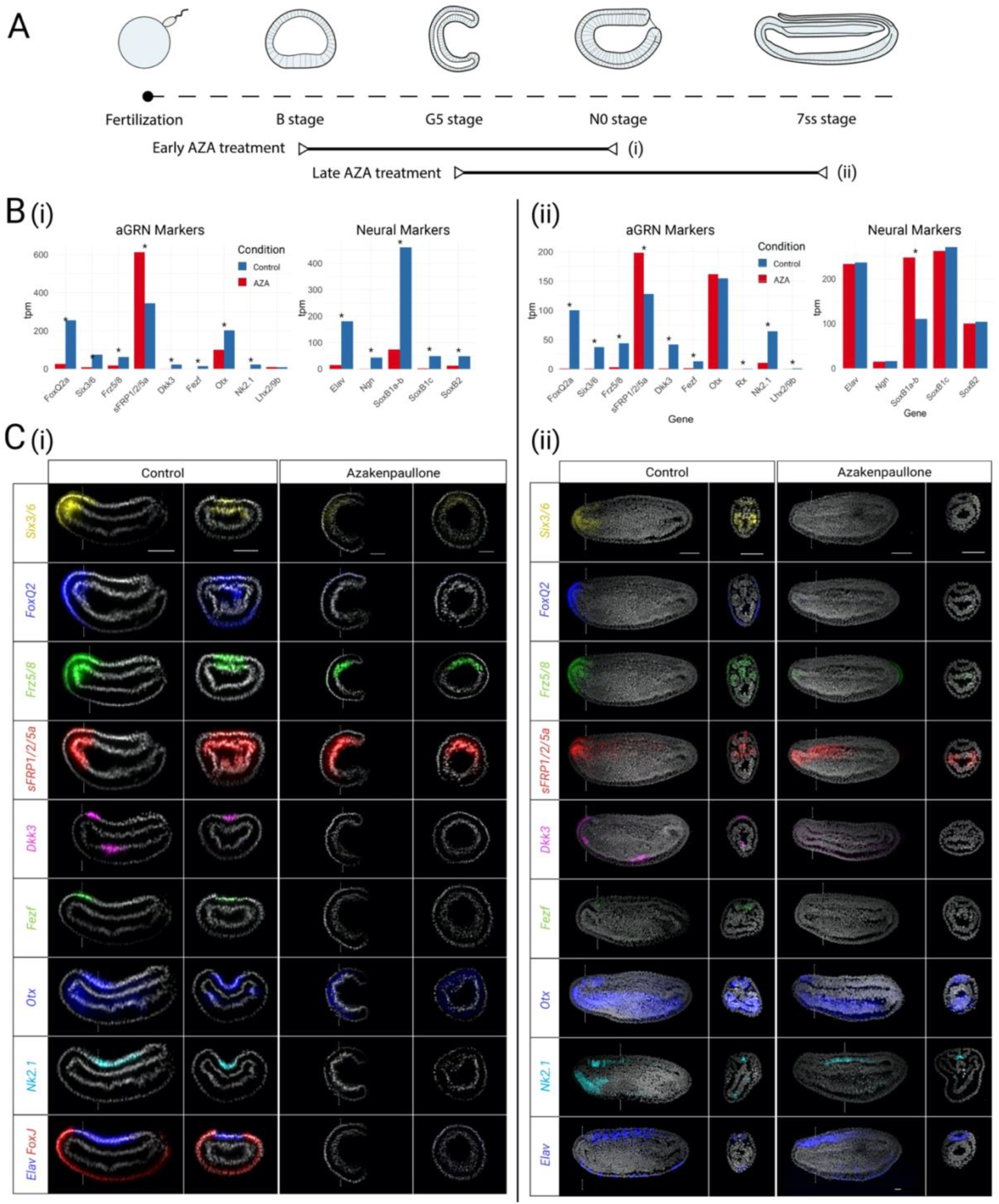
Characterization of Azakenpaullone-treated amphioxus embryos. **A** Experimental design of Wnt overactivation through early (B - N0 stage) and late (G5 – 7ss stage) Azakenpaullone (AZA) treatments. **B** Differential expression quantified by bulk RNAseq between control (blue) and AZA-treated (red) embryos in early (i) and late (ii) treatments. Asterisks indicate significant differences. **C** *In situ* HCR detection of aGRN markers *Six3/6* (yellow), *FoxQ2* (blue), *Frz5/8* (green), *sFRP1/2/5a* (red), *Dkk3* (magenta), *Fezf* (green), *Otx* (blue) and *Nk2*.*1* (cyan), of neural marker *Elav* (blue) and epidermal marker *FoxJ* (red) between control and early (i) or late (ii) AZA-treated embryos. While early Wnt overactivation causes loss of aGRN and neural plate, late treatments inhibit aGRN markers but not neural tube formation. Scalebars are 50µm.

Using RNA-seq, we first profiled all genes we found as part of the amphioxus aGRN (Figure 4B). Our results show that in early treated embryos all genes are noticeably downregulated following AZA treatment, with the sole exception of *sFRP1/2/5a*, which appears upregulated upon treatment (Figure 4Bi). We next investigated the context in which these changes were happening by looking at the expression of neural, ectodermal and mesodermal genes. We noticed that early pan-neural markers disappear following Wnt overactivation, consistent with previous published results shown by *in situ* hybridisation^83,84^, while mesodermal markers are upregulated (Figure 4Bi, Supplementary Figure 6A). Furthermore, we observed that genes marking the non-neural ectoderm at this stage also decrease significantly in expression in treated embryos, suggesting that the loss of neural tissue is not related to a failure of ectoderm progenitors to differentiate into neuroectoderm (Supplementary Figure 6A). To investigate this further, we also looked at the expression profile of amphioxus orthologs to known vertebrate markers for stem cells, finding upregulation after treatment, suggesting the ectoderm might remain in a naïve state in these embryos (Supplementary Figure 6A). In support of this, immunofluorescence for acetylated tubulin, shows that ectodermal cilia disappeared in treated embryos at the neurula stage (Supplementary Figure 6B). In late AZA treatments, most of the aGRN genes were strongly downregulated as for early treated embryos (Figure 4Bii). The exception was again *sFRP1/2/5a*, accompanied this time by *Otx*. In contrast with the early AZA treatment however, late treated embryos showed no significant decrease in neural or epidermal markers, while mesodermal genes were still upregulated (Figure 4Bii, Supplementary Figure 6A). Accordingly, detection of acetylated tubulin immunoreactivity revealed that both epidermal cilia and the cilia of the neural canal are visible in treated as well as control embryos (Supplementary Figure 6B).

Using *in-situ* hybridization, we next sought to spatially resolve the location of the genes identified as differentially expressed in our RNA-seq analysis (Figure 4C). Our *in-situ* profiling shows no trace of the aGRN in early treated embryos, as illustrated by an ectoderm-specific loss of *Six3/6, Frz5/8, sFRP1/2/5a* and *Otx*, which instead remained expressed in the mesendoderm, and a loss of *FoxQ2, Fezf* and *Dkk3* (Figure 4Ci). In these embryos, we also see the neural marker *Elav* and the early floor plate marker *Nk2*.*1* completely absent from the prospective dorsal neuroectoderm, indicating a loss of neural plate identity (Figure 4Ci). To rule out an off-target effect of AZA on dorsal-ventral specification, we co-profiled the dorsal determinant chordin (*Chd*) together with PSmad1/5/8, as a readout of ventral BMP activity, in normal and treated embryos. Our results show that whilst overactivation of the Wnt/β-catenin pathway clearly affects anterior-posterior patterning, it has no effect on dorsal-ventral specification at these early stages (Supplementary Figure 6B) and proves that the loss of neural plate identities in these embryos is not due to a loss of dorsal identity.

Our *in-situ* profiling in late treated embryos shows even more clearly that the aGRN is lost upon Wnt overactivation (Figure 4Cii). Again, *sFRP1/2/5a* and *Otx* remain strongly expressed in the mesendoderm, indicating the transcripts detected by RNA-seq profiling are mesendoderm specific. In contrast with the early treatment, and as predicted by our RNA-seq profiling, late treated embryos retain Elav expression in the neural plate (Figure 4Cii). We find this neural plate expresses *Nk2*.*1*, indicating that floor plate identities might be established at the latest at early stages of gastrulation. However, these late treated embryos have a continuous *Elav* domain with a rostral end coincident with the anterior neural border, clearly contrasting with that of control embryos, where *Elav* has a segmented pattern and is absent from the anterior neural border (Figure 4Cii). To investigate this further, we segmented the neural plate of late treated and control embryos using the IMARIS software (Figure 5A). Our analysis found no significant difference in the cell number (n=12, Student t-test, p=0.607) or volume (n=16, Student t-test, p=0.591) of the neural tissue in treated and control embryos (Figure 5A), suggesting the loss of aGRN markers is not due to a loss of anterior nervous tissue. In search of an alternative explanation, we set out to investigate the identity of the anterior nervous tissue in these late AZA treated embryos. To this end, we co-profiled *in-situ* the expression of the anterior markers *Otx* and *Pax4/6* together with *Gbx*, which marks the posterior border of the brain at the level of the first somite boundary, and *Hox1*, whose expression starts at the third somite within the putative hindbrain region^85^ (Figure 5B). Our results show that only a portion of cells in the posterior domain of *Otx* and *Pax4/6* remain in treated embryos. These cells are located at the anterior boundary with Gbx, which upon Wnt overactivation shifts to the anterior border of the neuroectoderm. This anterior shift in *Gbx* expression is accompanied posteriorly by an anterior shift in *Hox1* expression, indicating that the anterior brain, where the aGRN is acting, loses its anterior identity in late AZA treated embryos by acquiring a posterior identity.

**Figure 5.**
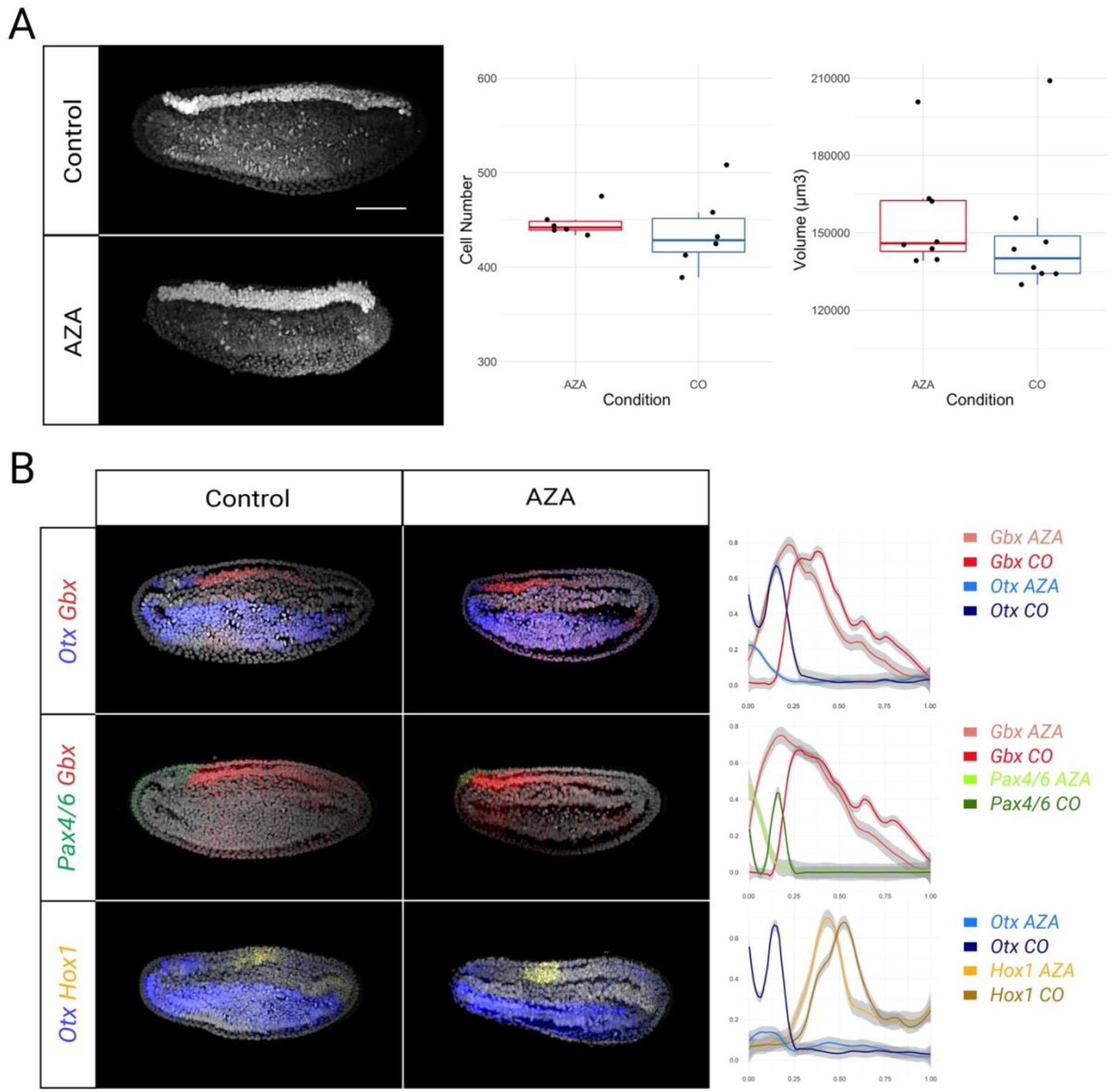
Loss of aGRN causes posteriorization of the anterior neural plate. **A** AZA treatment does not change the position, cell number and volume of the neural plate in 7ss stage embryos. Neural plate nuclei are false-colored in white. **B** Fate of anterior neural plate following Wnt overactivation: anterior markers *Otx* (blue) and *Pax4/*6 (green) restrict to the anteriormost tip of the neural plate, losing their anterior expression domain, while posterior markers *Gbx* (red) and Hox (yellow) expand anteriorly in AZA-treated embryos. Scalebars are 50µm.

Taken together, our data suggest that anterior aGRN genes in amphioxus are negatively regulated by Wnt signalling, similarly to ambulacrarian larvae, in which Wnt overactivation causes the loss of anterior ectodermal structures, both neural and non-neural. Furthermore, our results show that whilst late Wnt overactivation does not affect the number of cells specified to become neurons in the neural tube, the aGRN in amphioxus is essential for neurons to acquire an anterior brain identity.

### Apical patterning is essential to set the hypothalamic identity of neurons in the anterior brain of amphioxus

Having found that the aGRN in amphioxus is essential for the specification of ANE fate, we set out to investigate which anterior brain identities were particularly lost. Two lines of evidence pointed at the hypothalamus as the brain region potentially most affected by the loss of the aGRN. First, we found a strong signal for the aGRN to be operating during the specification of hypothalamic cells in the zebrafish brain (see Figure 1). Second, we found that *Nk2*.*1* and *Fezf*, two early markers of hypothalamic fate in vertebrates (see supplementary Figure 1F)^86^, are lost from the amphioxus anterior CNS upon aGRN silencing by Wnt overactivation (see Figure 5). In normal development, the posterior limit of *Fezf* would meet the anterior limit of *Nkx2*.*1* expression at the intercalated *Six3/6*-negative region, which we recently defined as hypothalamic for co-expressing the vertebrate hypothalamic markers *FoxD* and *Otp*^77^. Given this, we reasoned that the loss of *Six3/6* expression in AZA treated embryos might imply a loss of the hypothalamic brain.

To test this hypothesis, we built up on our previously published *Otp, FoxD* and *Six3/6* dataset by co-analyzing the expression in early amphioxus larvae of additional key hypothalamic markers, such as *Bsx*^87^ and *Hmx* (pro-orthologous of *Hmx2* and *Hmx3*)^88^, which are expressed in zebrafish hypothalamic cells (Figure 6A and Supplementary Figure 7A). We found that *Bsx* at the 12ss and 14ss co-localizes with *Otp* in cells bilaterally arranged at the periphery of the preinfundibular brain, which lies in the *Six3/6-/FoxD+/Otp+* domain that we define as hypothalamic (Figure 6A). Complementing this *Bsx* pattern, we detect *Hmx* expression in anterior pre-infundibular cells, again within the Six*3/6-/FoxD+/Otp+* domain (Figure 6Aii). In line with this being part of the amphioxus hypothalamus, we find the middle *Bsx* domain absent from embryos in which cell division has been arrested from the 7ss (Supplementary Figure 7B). These results recapitulate and expand what we already reported for *Otp*^77^, corroborating that the amphioxus hypothalamus develops by intercalated growth in neurulating embryos and demonstrating that *Bsx* positive cells in the Six*3/6-/FoxD+/Otp+* domain are part of this developmental program (Figure 6C).

**Figure 6.**
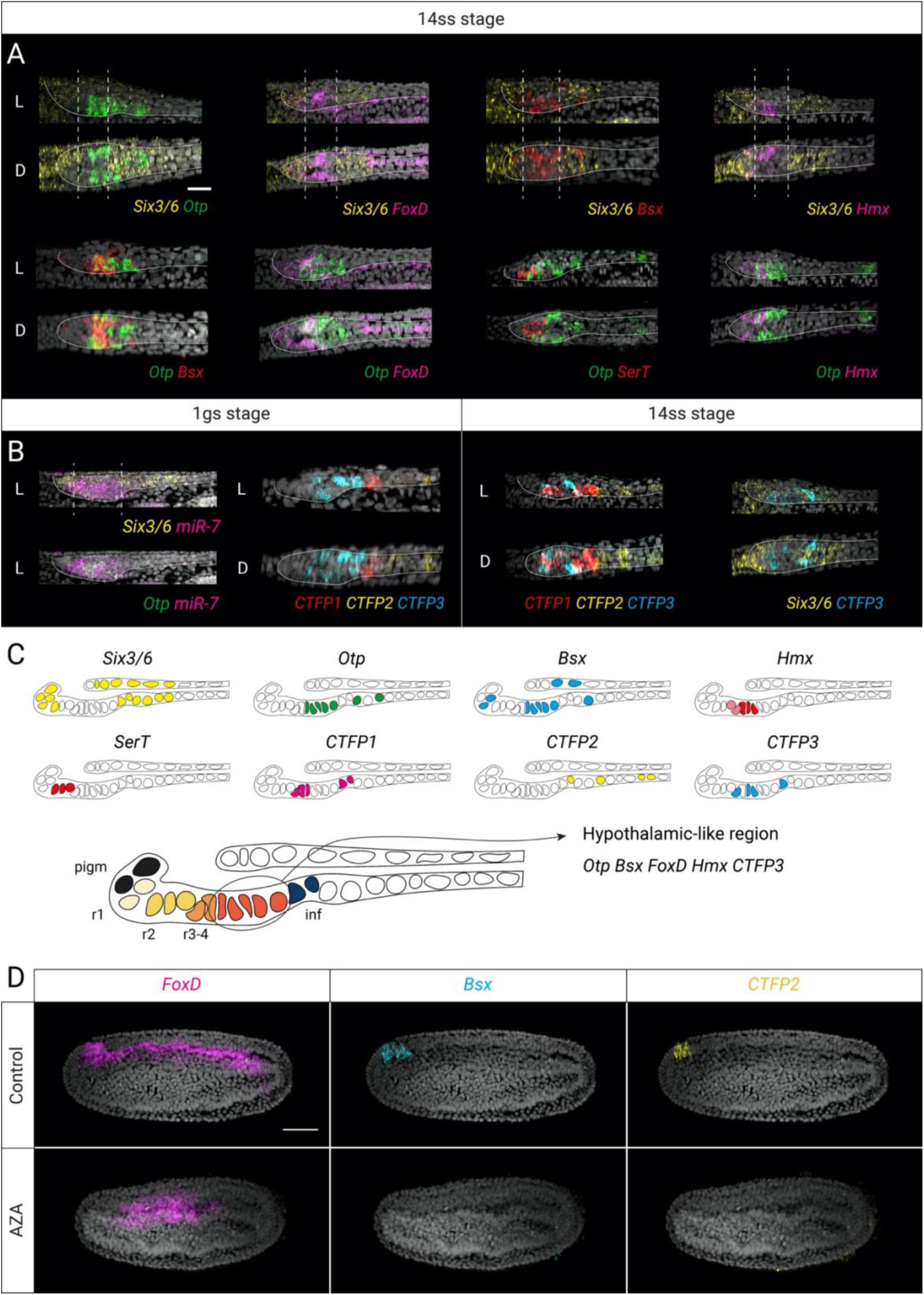
Molecular characterization of a hypothalamic-like region in the anterior brain of amphioxus larvae. **A** Co-detection of *Six3/6* (yellow) and hypothalamic markers *Otp* (green), *FoxD* (magenta), *Bsx* (red) and *Hmx* (magenta) in the cerebral vesicle of early larvae (14ss) in lateral (L) and dorsal (D) view. *SerT* (red) is used as a reference for the position of serotonergic neurons. **B** Localization of *miR-7* mature transcripts (magenta) related to expression of *Six3/6* (yellow) and *Otp* (green) in lateral views of the cerebral vesicle of 1 gill slit (gs) larvae, and expression of the three calcitonin family peptides in lateral and dorsal views of the cerebral vesicle in 1gs and 14ss larvae. **C** Schematic overview of gene expression and cell type distribution in the anterior cerebral vesicle of amphioxus. The circle highlights the hypothalamic-like region. **D** Loss of early expression of hypothalamic markers *Fox, Bsx* and *CTFP2* in the anterior neural plate of late Azakenpaullone-treated embryos. Scalebars are 20µm in A and 50µm in D.

The vertebrate hypothalamus is also characterized by the expression of the microRNA miR-7, which in addition is associated with neurosecretory centers in protostomes and deuterostomes^89^. In amphioxus, miR-7 has been reported in the brain and the pre-oral pit^90^, but its precise relation to the *Six3/6*-/*FoxD*+/*Otp*+ hypothalamic region is unknown. Given this, we co-profiled the expression of miR-7 in 1gs larvae together with *Otp* and *Six3/6* and found that while mir-7 is broadly expressed in the ventrolateral cerebral vesicle, the entire pre-infundibular *Otp* cluster is mir-7 positive (Figure 6B). This prompted us to look further into putative neurosecretory centers in the amphioxus brain. We investigated the distribution of the three amphioxus neuropeptides of the calcitonin/CGRP family (calcitonin family proteins, CTFPs)^91,92^. In 1gs larvae, *CTFP3* is indeed expressed in the anterior cerebral vesicle within the miR-7 domain (Figure 6B), while *CTFP1* is restricted to the post-infundibular region and *CTFP2* is expressed more posteriorly. By looking at the developmental distribution of the three paralogs we highlighted a dynamic expression pattern: *CTFP1* and 2 are first co-expressed in the anterior nervous system at the 7ss stage (Supplementary Figure 5). While *CTFP1* continues to be expressed in both the anterior and posterior cerebral vesicle, mostly in medial cells, *CTFP2* abruptly changes its pattern to mark cells in the posterior cerebral vesicle and the trunk region of the nervous system at 14ss (Figure 6B). At the same stage, *CTFP3* appears in the cerebral vesicle and the infundibular organ region (Figure 6B).

Having found that the preinfundibular brain of amphioxus has a strong hypothalamic identity and neurosecretory capacity, we then investigated whether such identities are maintained in the absence of the aGRN. To this aim we co-profiled *FoxD, Bsx, CTFP1* and *CTFP2*, which are already expressed at the 7ss stage, in late AZA treated embryos, either *in situ* or by RNA-seq (Figure 6D and Supplementary Figure 7D). Our results show no trace of hypothalamic or neurosecretory identities in the absence of the aGRN by Wnt overactivation, showing that the hypothalamic identity is lost following the loss of the aGRN caused by Wnt overactivation.

## Discussion

AOs have long been at the centre of the debate on the evolution of nervous systems. The reason for their evolutionary relevance is that AOs are present in the ciliated primary larvae of a number of taxa across metazoans, including cnidarians, spiralian protostomes and ambulacrarian deuterostomes (Echinodermata + Hemichordata)^12,13,20,93^. AOs are therefore one of the few unifying features across distantly related animals with the most diverse neural architectures, ranging from nerve nets to ventrally centralized nerve cords^12,14,17,22,24,28,94–97^. Given this, the absence of AOs in chordates has given rise to numerous hypotheses aiming to explain the loss of AOs and gain of a dorsally centralized neural tube in this group of animals. Perhaps the most extreme was that proposed by Garstang, postulating that the AO was internalized in the chordate lineage by the fusion of lateral ciliary bands and became incorporated into the vertebrate brain^98^. This idea has found little support from modern studies, but the notion that the AO is somehow evolutionarily connected to the origin of the vertebrate forebrain has never been abandoned^25,99,100^.

In the wake of the genomic era, it has become clear that AOs are specified by genes whose orthologous counterparts are expressed in the anterior CNS of vertebrates, and so the parallel between AOs and the vertebrate brain has been proposed a number of times^25,45,56,101,102^. However, a direct connection between these two different neural organizations has never been proved. Our work solves that puzzle by identifying in amphioxus an aGRN with a highly similar gene composition and expression dynamics to that specifying the AO of sea urchins. We show that in amphioxus the aGRN is active from early development and is essential to define the anterior part of the nervous system as forebrain and to establish the identity of retinal and hypothalamic cell types. By exploiting single-cell genomics approaches, we find the same aGRN also acting during the development of the zebrafish hypothalamus and retina, demonstrating that amphioxus is not the only chordate utilizing this aGRN for developing specific parts of the forebrain, therefore suggesting that this is a conserved feature of chordates.

Our findings are supported by the combinatorial analysis of 22 genes through amphioxus development using RNA-Seq profiling and multiplexed *in situ* hybridization. Most importantly, we identify individual cells with an active aGRN across invertebrate and vertebrate deuterostomes: in an ambulacrarian (sea urchin), a basally branching chordate (amphioxus) and a vertebrate (zebrafish), thus establishing a link between AOs and the chordate ANE. Our work also addresses some of the challenges posed by the “chimeric brain hypothesis”^56^ and the earlier “animal-axial hypothesis”^57^, namely the conflicting fate of ANE precursors in invertebrates and chordates. In invertebrates the ANE gives rise to the AO and is specified within an AP (defined by the early expression of *Six3/6* and *FoxQ2* from early development^11^) that comprises both epidermal and neuroectodermal derivatives^45^. Conversely, in vertebrates the ANE does not form within a recognizable AP, but is specified within the neural plate and gives rise to forebrain identities only^47,59^. Our results show that the amphioxus ANE forms within an early AP expressing *Six3/6* and *FoxQ2* and gives rise to both AO and forebrain identities. We see early AP genes (EAP: *FoxQ2, Six3/6, Frz5/8, sFRP1/2/5a, Dkk3* and *Otx*) active in anterior ectodermal and neuroectodermal progenitors, as shown by co-profiling with *Ngn, Elav* (neural markers) and *FoxJ* (non-neural ectoderm marker) (see Figure 2). Late AO genes (LAP: *Fezf, Rx, Nk2*.*1*, and *Lhx2/9*, including *SerT*) are instead primarily active in precursors that are already committed to a neuronal fate (see Figure 3). Multiplexed profiling at later developmental stages reveals these committed neuronal precursors to become part of the anterior photoreceptive complex and neurosecretory center of the amphioxus brain. This result is consistent with the hypothetical contribution of the ancestral apical brain proposed under the chimeric brain evolutionary scenario^56^. In support of this, we find that the serotonergic cells of the frontal eye complex co-express *Fezf* (a LAP gene) as in sea urchin; and that the hypothalamic region, defined as the *Six3/6-/FoxD+/Otp+* region of the brain, expresses *Bsx* and *Hmx*, together with *mir-7* and two of the three amphioxus calcitonin genes, *CTFP1* and *CTFP3*, indicating its neurosecretory nature. Most importantly, we see precursors of all these cell types missing upon GSK3 inhibition, demonstrating that the aGRN specifying retinal and hypothalamic cell identities in amphioxus depends on Wnt signalling.

Our functional assays further show that the biphasic specification of the apical (anterior) ectoderm in amphioxus is regulated by Wnt signalling, as previously shown in sea urchin in a series of publications by the Range lab^31,36,43^. In doing so, we describe a new role of Wnt signalling in amphioxus, which adds to the previously reported function of early Wnt/β-catenin signalling in the specification of dorsal fates and of zygotic Wnt in ventral/posterior fates^83,84,103^. We see that *FoxQ2* and *Wnt8* are expressed in amphioxus from the blastula stage in two mutually exclusive domains, at the animal and vegetal side of the embryo respectively. This early expression greatly resembles the one seen in ambulacrarian larvae, and has no counterpart in vertebrates, as *FoxQ2* has not been detected in any vertebrate embryos before organogenesis and the gene is lost in placental mammals^51^. Moreover, at this stage the expression is radial and does not show any type of DV regionalization, suggesting that it is not directly influenced by Nodal signalling acting at the posterior-dorsal side at this stage^84^. The radial distribution continues during early gastrulation, when a band of dorsal neural progenitors appears within the *FoxQ2* domain. Only around the end of gastrulation, *FoxQ2* transcripts disappear from the ventral side of the embryo and starts to restrict towards the anterior-dorsal tip during neurulation, together with *Six3/6* and *Frz5/8*. During the restriction, two secreted Wnt inhibitors, *sFRP1/2/5a* and *Dkk3*, are produced from the anterior-dorsal tip of the embryo. This suggest that Wnt signalling acts through mutually repressive interactions to restrict the size and position of the aGRN to the dorsal anterior (apical) side of the embryo, as described in ambulacrarians^31,36,58^. This territorial restriction happens exclusively in the ectoderm, since GSK3 inhibition does not block the mesodermal expression of any of the EAP genes.

The suppression of the aGRN by Wnt overactivation was observed throughout gastrulation and neurulation, but the treatment had different effects on the specification of the neuroectoderm depending on developmental time. While Wnt overactivation from the blastula stage caused the loss of neural plate, treatment during neurulation did not affect the size and position of the neuroectoderm but led to the loss of anterior (forebrain) fate. Taking into consideration that Wnt overactivation does not result in a loss of dorsal identity, as revealed by the similar *Chd* and Smad distribution in control and treated embryos, we propose that early zygotic Wnt signalling is key to provide positional information to the ANE, and in doing so it is essential to set the position of the brain through aGRN-mediated anteriorization signals. Little is known about the mechanism by which the position of the brain is set in vertebrates, although a requirement from the rostral non-neural ectoderm to form the ANR (anterior neural ridge) has been observed^55^. It is still unclear if Wnt signalling is involved in setting the ANR itself, but it has been reported that in the zebrafish the ANB is a source of SFRPs that act to repress posteriorizing Wnt signals well before the formation of the ANR, which is essential to form the brain^47,59,104^. A series of studies from the Weinberg lab^105–107^ have also shown that in zebrafish the AP patterning of the neuroectoderm is not dependent on the primary organizer (which is positioned by maternal β-catenin), but it is affected by zygotic Wnt signalling. Given this, our results suggest that a similar aGRN as the one identified in amphioxus, but devoid of *FoxQ2*, might be also operating in vertebrates to position the forebrain early in development.

Taking all results in consideration, we propose that the aGRN is an ancient signalling system, also conserved in chordates, that specifies anterior sensory-neurosecretory neural fate (Supplementary Figure 8). We have recently shown that the serotonergic neurons of the amphioxus frontal eye and major parts of the hypothalamus develop by addition of new cellular material during neurulation, as opposed to the previous notion that cells in this area would transform into those precise identities^77^. Hence, this suggests a scenario in which newborn cells during neurulation would be immediately under the influence of the aGRN, thereby coupling an ancient signalling system to the revolutionary developmental innovation which constitutes forming a dorsal neural tube through neurulation in chordates.

## Materials & Methods

### scRNA-seq analysis

#### Sea urchin developmental dataset

Normalized single-cell RNA-sequencing data from the gastrulating *Strongylocentrotus purpuratus* was obtained from^63^ (GEO accession GSE149221). Batch correction was first performed using *fastMNN* (using 20 nearest neighbors), implemented in the *batchelor* R package^108^ on the top 2000 highly variable genes. This gene set was obtained by using the *‘modelGeneVar’* and ‘*getTopHVGs’* functions in the *scran* R package^109^, which accounts for the mean-variance relationship of each gene. Samples were then integrated in reverse order of their developmental stage so that samples are merged with decreasing numbers of cells. The 50 corrected principal components were subsequently used to obtain a reduced dimensionality UMAP. This was performed using the UMAP implementation in Scanpy^110^ using 75 nearest neighbors and a 0.9 minimum distance parameter.

##### Reannotation

The Foster et al. 2020 cell type annotations were obtained by cross-referencing clusters in the raw Seurat object with labels given in the original publication. We then performed a finer annotation of apical organ cell types. By averaging the expression of apical patterning genes across developmental stages, we found a clear distinction in the genes expressed from the 8-cell to early blastula stages and from the hatched blastula to late gastrula stages (Supplementary Figure 1C). This motivated the separation into an ‘early’ and ‘late’ apical plate cell type defined by those apical patterning genes expressed at early and late timepoints. To classify cells of the apical plate, the sea urchin dataset was iteratively clustered and filtered based on the expression of these apical plate cell type markers. Leiden clustering, implemented in Scanpy^111^, was performed at various resolutions (resolution parameter ranging from 1 to 8) on the 50 corrected principal components. These clusters were then assigned a cell type by comparing with the expression profiles of apical patterning genes, the serotonergic neuron marker, *Tph*, as well as the developmental stages and original Foster 2020 annotations.

#### Zebrafish - developmental dataset

Raw counts and metadata were obtained from^67^. These raw counts were normalized and log-transformed using the *‘computeSumFactors’* and *‘logNormCounts’* functions from the *scran* R package. The two-dimensional, force-directed graph representation in Figure 1Bi, Ci was replicated from the original publication using the provided Matlab script (https://github.com/wagnerde/STITCH/) and graph visualization software, *Gephi*^112^. In this setup, Wagner et al. link cells across developmental stages using a novel graph construction algorithm, STITCH. This graph is then visualized using the ForceAtlas2 algorithm^113^.

##### Reannotation

Cells annotated by Wagner et al. as ‘diencephalon’ from stages 14hpf, 18hpf and 24hpf were isolated to obtain a more refined annotation according to the prosomeric model. As with the sea urchin, multiple rounds of Leiden clustering and filtering was performed. The STITCH graph was used for both Leiden clustering and ForceAtlas2 visualization with Scanpy (Supplementary Figure 1E,F). Hypothalamus cells were then separated from the diencephalon according to the expression of hypothalamus (*fezf2, nkx2*.*1,rx3*) and posterior diencephalon (*barh2,pitx2, pitx3*) markers, reported in previous studies.

#### Amphioxus developmental dataset

Normalized single-nucleus RNA-seq data was obtained from^73^ via links provided in the accompanying shiny app (https://lifeomics.shinyapps.io/shinyappmulti/). Cell type and stage annotations were provided in the downloaded Seurat object, as well as UMAP embedding coordinates. Annotations for each amphioxus gene ID were obtained from the original study (available through the link above and via https://github.com/XingyanLiu/AmphioxusAnalysis/tree/main/resources). However, the gene IDs used for ‘*Dkk3*’ and ‘*Dkk1*’ was confirmed using BLAST best alignment hits. UMAP expression plots were created using Scanpy.

#### Stage heatmaps

The sea urchin, amphioxus and zebrafish datasets were filtered to contain only ectoderm-related cell types. Specifically, cell types annotated as “SN”, “Late AP”, “Early AP”, “Other ectoderm” and “Other neural” were retained in the Foster et al. 2020 sea urchin dataset. The Ma et al. 2022 amphioxus dataset was filtered based on the lineage annotations provided in the original study. Specifically, “B_0”, “B_1”, “Epithelial ectoderm” and “Neural ectoderm” cells were retained. Similarly, only “Forebrain / Optic”, “Midbrain”, “Hindbrain / Spinal Cord”, “Neural Crest” and “Epidermal” cell types were used from the Wagner et al. 2018 zebrafish dataset. After filtering, normalized gene expression values were averaged across cells within each developmental stage and min-max normalized within each gene. This was then plotted using the *ComplexHeatmap* package.

#### Zebrafish - hypothalamic dataset

Single-cell RNA-seqeuncing data from^86^ was also used to query later stages of zebrafish brain development. Processed Seurat objects from each developmental stage were downloaded from GEO accession GSE158142 and combined into a single SingleCellExperiment object. The counts data were renormalised using the *‘multibacthNorm’* function from the batchelor R package, which removes systematic differences in coverage across batches. All batches in the combined dataset were then integrated together using *fastMNN* with 20 mutually nearest neighbours. Batches were merged in reverse order of developmental stage and in order of decreasing numbers of cells. As in the analysis of the Wagner et al. 2018 dataset, the corrected principal components were used to compute a reduced dimensionality UMAP (15 nearest neighbours, 0.5 minimum distance). The dataset was then filtered to include only those cells annotated by the original authors as being related to the hypothalamus (labelled as *’hypothalamus/telencephalon, gabaergic’, ‘hypothalamus’, ‘hypothalamus’, ‘ventral diencephalon/hypothalamus’, ‘hypothalamus neurons’ or ‘Hypothalamus’*).

### Amphioxus husbandry, spawning and embryo fixation

Adult individuals of European amphioxus *B. lanceolatum* were collected from the sand in Banyuls-sur-Mer, France, and transported to Cambridge, UK, where they were kept in a custom-made facility at the Department of Zoology. Here amphioxus were maintained and gametes obtained as described in^114^. After *in vitro* fertilization, embryos were raised in Petri dishes in filtered artificial salt water (ASW) in an incubator at 21°C. For embryos at 4 dpf, from the 48 hours stage embryos were fed with a mix of algae. At the desired stage embryos were collected and fixed in ice-cold 3.7% Paraformaldehyde (PFA) + 3-(N-morpholino) propane sulfonic acid (MOPS) buffer for 12 hours, then washed in sodium phosphate buffer saline (NPBS), dehydrated and stored in 100% methanol (MeOH) at -20°C. For RNA sequencing, live embryos were transferred in a RNAse-free Eppendorf, fast-frozen with liquid nitrogen and then stored at -80°C before processing. The nomenclature for early stages is taken from^115^, while during neurulation the number of somites is used to unbiasedly identify the embryonic stage.

### Amphioxus RNAseq analysis

#### Amphioxus developmental dataset

The *B. lanceolatum* transcriptome and genome described in^74^ were downloaded via the link provided. Additionally, the FASTA files for the mitochondrial *B. lanceolatum* genome and transcriptome sequences, as described in^116^, were retrieved from NCBI (AB19483). The four references were used to generate a reference index using Salmon (version 1.3.0)^117^.

Raw read quality was assessed using FastQC (version 0.11.9) and adapter contamination was removed by trimming the reads using Trimmomatic version 0.39^118^. Transcript expression was quantified with Salmon (version 1.3.0). Transcript level expression values were imported into R and summarised to gene level using the Bioconductor package _tximport^119^. Counts were then normalized for library size and composition bias using DESeq2^120^.

#### Differential expression of Azakenpaullone treated embryos

Control and treated embryos frozen at -80°C were ground with an electric pestle in a 1.5ml tube while thawing and then total RNA was extracted using Norgen Total RNA Purification Plus Kit (Catalog n° #48300). Purified RNA was stored in -80°C. RNA quantity and quality was checked using a BioAnalyzer. Libraries were prepared using the Illumina TruSeq Stranded mRNA kit (#20020595) following the manufacturer instructions and sequenced on a NovaSeq 6000 system.

Raw read quality was assessed using FastQC (version 0.11.9) and reads were trimmed using fastp version 0.21.0^121^ to remove adapter contamination and poly-G tails. Transcript expression was quantified with Salmon (version 1.3.0). Transcript level expression values were imported into R and summarized to gene level using the Bioconductor package _tximport^119^. Differential expression analysis between AZA treated and Control groups was carried out using DESeq2^120^, the “local” fit method was used for dispersion estimates and multiple testing correction was carried out using the Benjamini-Hochberg method^122^.

### Drug treatments

Fertilized eggs were raised at 21°C. At 4.5 hpf (blastula stage) embryos were transferred into small beakers with 10ml of 10µM Azakenpaullone (AZA) or dimethylsulfoxide (DMSO as a control) in filtered ASW and left to develop at 21°C up to 12 hpf (early neurula stage) or 21 hpf (7 ss stage). AZA is an inhibitor of GSK3β and has been used in marine larvae in previous works^11,82^. At 12 hpf the embryos were either fixed in 3.7% PFA in MOPS buffer and stored in methanol at -20°C or frozen in liquid nitrogen and stored at -80°C. For all the experiments, three replicas for treatment and control were used.

### Probe synthesis and *In situ* hybridization chain reaction (HCR)

Sequences for the genes used in this study were reconstructed through the Reciprocal Best Hits approach: genes from other amphioxus and deuterostome species were used to identify candidate orthologs in the *B. lanceolatum* transcriptome^74^; the top candidates were then compared with published transcriptomes using BLAST. Reconstructed sequences were then sent to Molecular Instruments, Inc. to prepare probe pairs used in version 3 *in situ* HCR, performed on embryos as described in^76^: briefly, samples were rehydrated in NPBS + 0.1% Triton X, (NPBT) bleached for 30 minutes with a solution of 5% Deionized formamide, 1.5% H2O2, 0.2% SSC in nuclease-free water and permeabilized for 3 hours in NPBS + 1% DMSO + 1% Triton. The embryos were incubated in Hybridization Buffer (HB, Molecular Instruments) for 2 hours and then the probes were added in HB overnight at 37°C. The following day probe excess was removed with Wash Buffer (Molecular Instrument) followed by 5X-SSC + 0.1% Triton X. The samples were then incubated in Amplification Buffer (AB, Molecular Instruments) for 30 minutes and then left overnight in the dark at room temperature in AB + 0.03µM of each hairpin (Molecular Instruments). Embryos were washed in the dark in 5X-SSC + 0.1% Triton X and incubated overnight with 1 μg/mL DAPI in NPBT, then washed in NPBT and transferred in a glass-bottomed dish in 100% glycerol. Imaging was performed on an Olympus V3000 inverted laser scanning confocal microscope.

### Immunohistochemistry

Fixed embryos were processed for immunofluorescence as described in^77^. Briefly, embryos were rehydrated and bleached as described above for *in situ* HCR and permeabilized overnight in NPBS + 1% DMSO + 1% Triton. The following day, samples were blocked in blocking solution (NPBS + 0.1% Triton + 0.1% BSA + 5% NGS) for 3 hours and then incubated overnight at 4°C in block with primary antibodies: mouse anti-acetylated tubulin (Sigma, T6793) at 1:250 or rabbit anti-phosphorylated Smad1/5/8 (Cell Signaling, 9511S) at 1:100. The following day embryos were thoroughly washed with NPBS + 0.1% Triton and incubated first in blocking solution for 3 hours and then overnight at 4°C in the dark in blocking solution with DAPI (1:500) and secondary antibodies: goat anti-mouse (Invitrogen 84540) and goat anti-rabbit (Abcam 150083), both at a dilution of 1:250. Samples were finally washed in NPBS + 0.1% Triton, mounted in 80% glycerol and imaged with on an Olympus V3000 inverted laser scanning confocal microscope.

### Image analysis

Z-stacks of 7ss embryos treated with AZA or DMSO and stained with HCR were imported in IMARIS (IMARIS 9.7.2, Bitplane, Oxford Instruments). The neural plate was manually segmented in 16 embryos (8 AZA treated, 8 controls) by drawing splines around the cell nuclei visible through the DAPI signal every 4 slices in transverse section. At this embryonic stage, the neural plate is clearly distinguishable from the rest of the ectoderm in cross section; in addition, *Elav* signal was used in half of the samples to validate the precision of the segmentation. In the segmented neural plate cells were counted manually using the DAPI signal, while volumes were calculated automatically by IMARIS.

## Data and code availability

All code used to replicate the single-cell RNA-seq analysis and figures can be obtained at https://github.com/ebglab/Amphi-Apical-Organ/. External datasets used are available via links provided in the methods.

## Acknowledgments

The authors would like to thank Matt Wayland from the Department of Zoology for his assistance during imaging, Katarzyna Kania from the Genomics Core Facility at CRUK Cambridge Institute for her help with RNA sequencing, Fadwa Joud and the Light Microscopy Facility at CRUK Cambridge Institute for assistance in the use of the IMARIS software and John Marioni and Toby Andrews for the feedback and discussions during the project and the writing of the manuscript.

## Author’s contributions

GG and EBG designed the project; GG and EBG designed experiments, performed embryo collection and drug treatments; GG performed *in situ* hybridization chain reactions, immunohistochemistry and RNA extraction for RNAseq; GG imaged embryos and performed image analysis; DK analysed scRNAseq datasets; AS and GG analysed RNAseq data; GG and DK wrote the original draft of the manuscript; GG, DK and EBG reviewed and edited the manuscript; EBG supervised the project and managed funding. All authors discussed the results and agreed to the submitted version of the manuscript.

## Funding

We acknowledge funding from the Whitten Trust to GG, and the Welcome Trust Mathematical Genomics Medicine Programme at the University of Cambridge (PFZH/158 RG92770) to DK. Work in the EBG lab was supported by the CRUK (C9545/A29580).

## Conflict of Interests

EBG has been an employee of Genentech since September 2022.

## Materials & Correspondence

Elia Benito-Gutiérrez

## Notes

### Competing Interest Statement

Elia Benito-Gutierrez has been an employee of Genentech since September 2022.

