## Supplementary Information for "An ancient gene regulatory network sets the position of the forebrain in chordates"

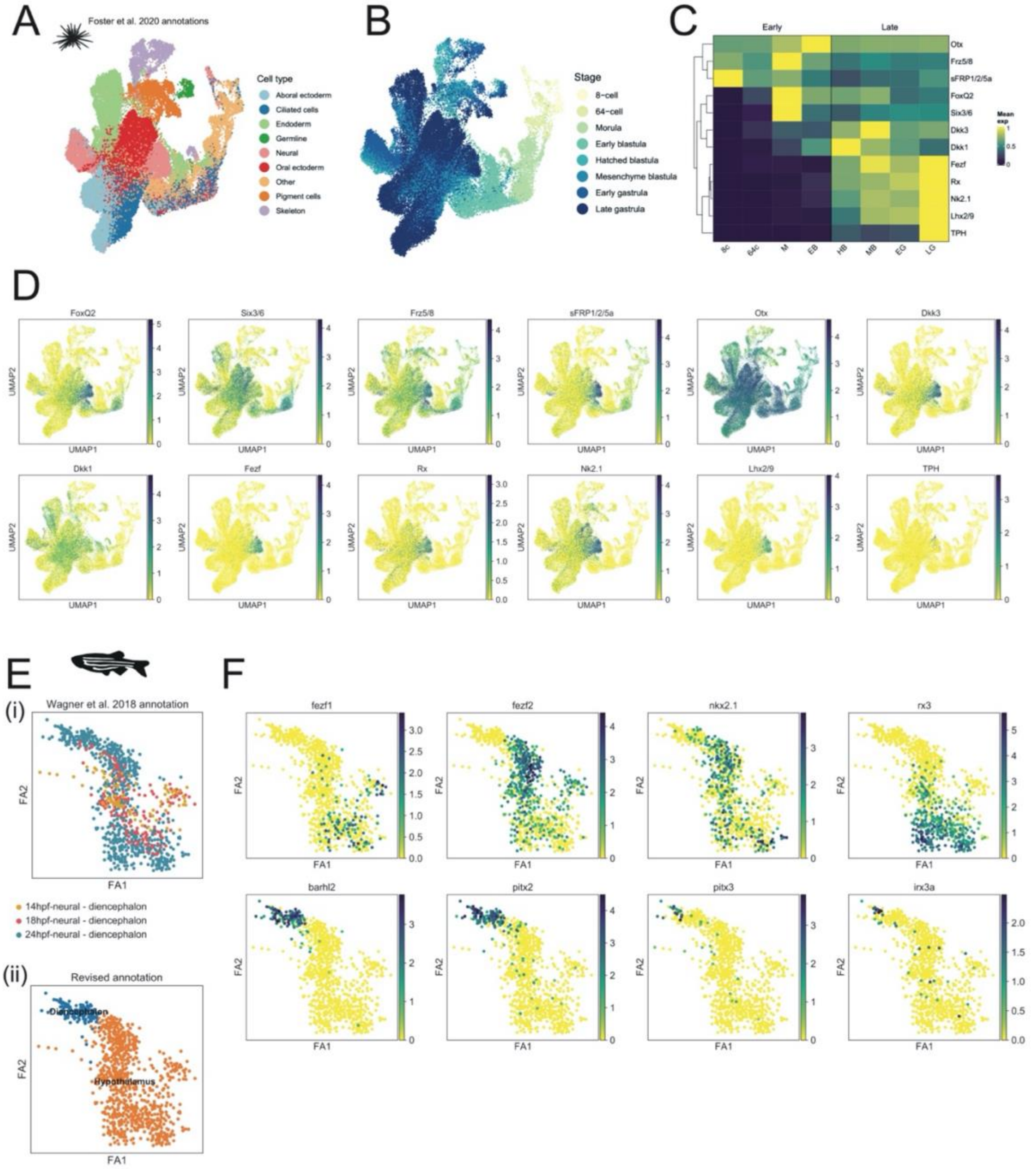


**Supplementary Figure 1. Re-annotation of sea urchin and zebrafish single-cell datasets.**

**A** UMAP plot of sea urchin single-cell dataset colored according to the original Foster et al. 2020 cell type annotations and **B** developmental stages. **C** Mean expression of apical patterning genes across developmental stages show two phases of expression representing early and late developmental stages. Values are min-max normalized across each row. The dendrogram and order of genes reflect hierarchical clustering across rows. 8c, 8-cell stage; 64c, 64-cell stage; M, morula; EB, early blastula; HB, hatched blastula; MB, mesenchyme blastula; EG, early gastrula; LG, late gastrula. **D** The expression of apical patterning genes, used to annotate apical organ cell types, across the entire sea urchin dataset. **E** ForceAtlas2 representation of Wagner et al. 2018 annotated diencephalon cells colored by **i)** the original annotation/developmental stage and **ii)** our revised annotation. **F** Expression of genes used to separate the diencephalon from hypothalamus.

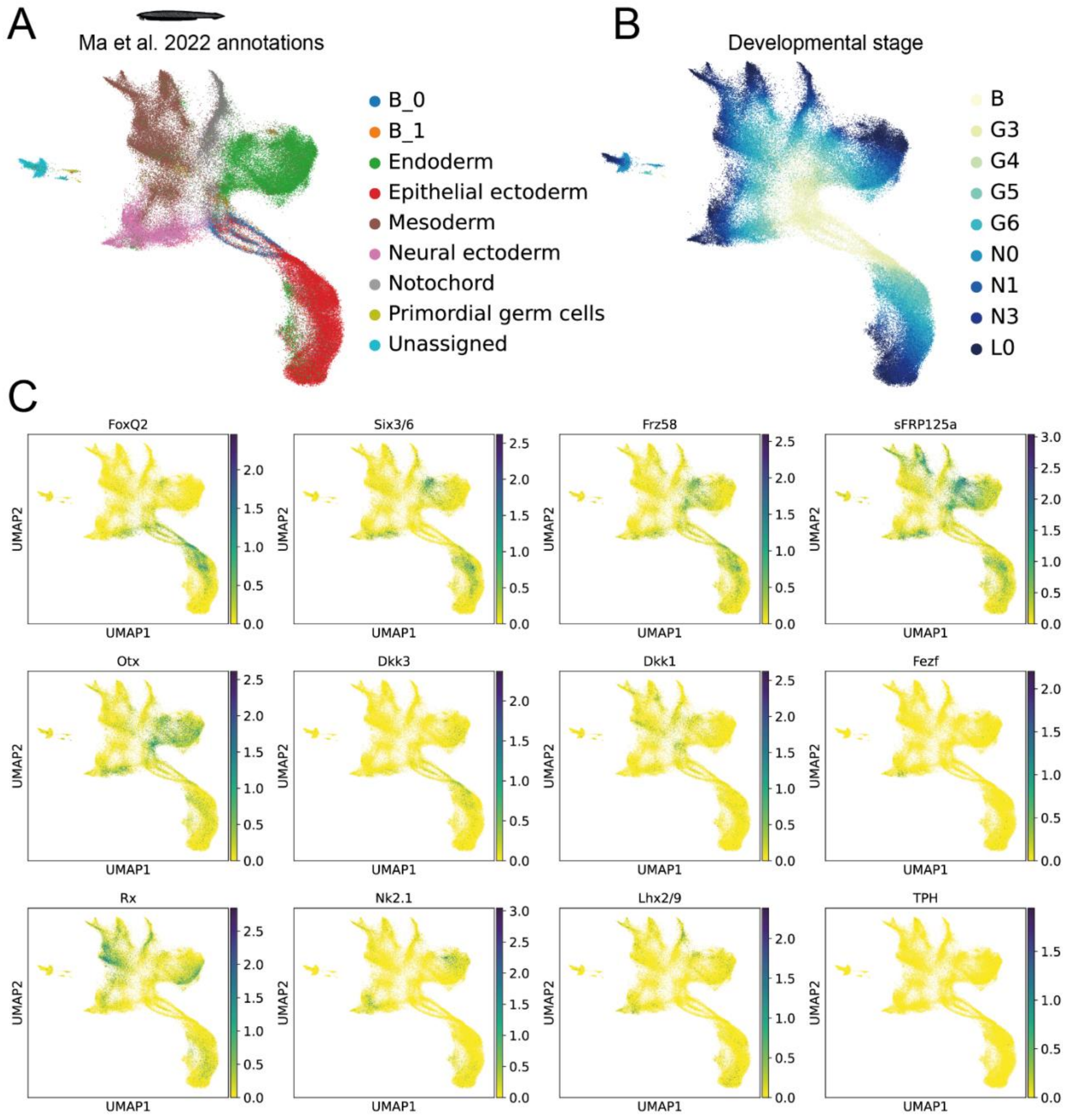


**Supplementary Figure 2.**

**A-B** UMAP plot of Ma et al. 2022 amphioxus snRNA-seq data colored by **A** lineage annotations and **B** developmental stages, provided in the original study. **C** Normalized expression of apical patterning genes visualized across the UMAP embedding. B: blastula; G3: early gastrula - G6: late gastrula; N1: early neurula – N3: mid neurula; L0: larval stage.

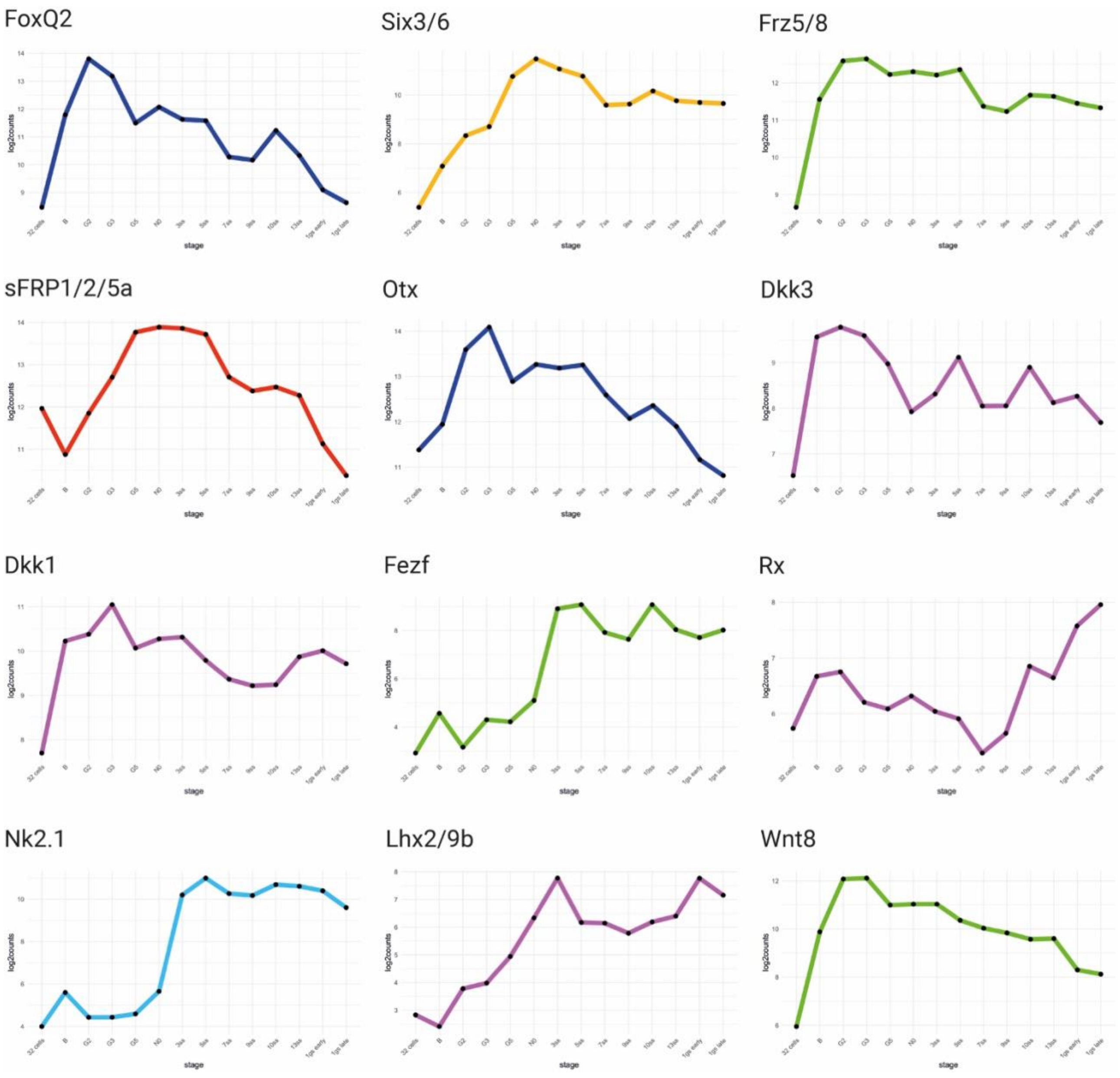


**Supplementary Figure 3. Differential expression of aGRN markers through amphioxus development.** Analysis performed on the bulk RNAseq dataset previously published by Marlétaz et al., 2018 for *B. lanceolatum* showing the progressive activation of early and late aGRN markers.

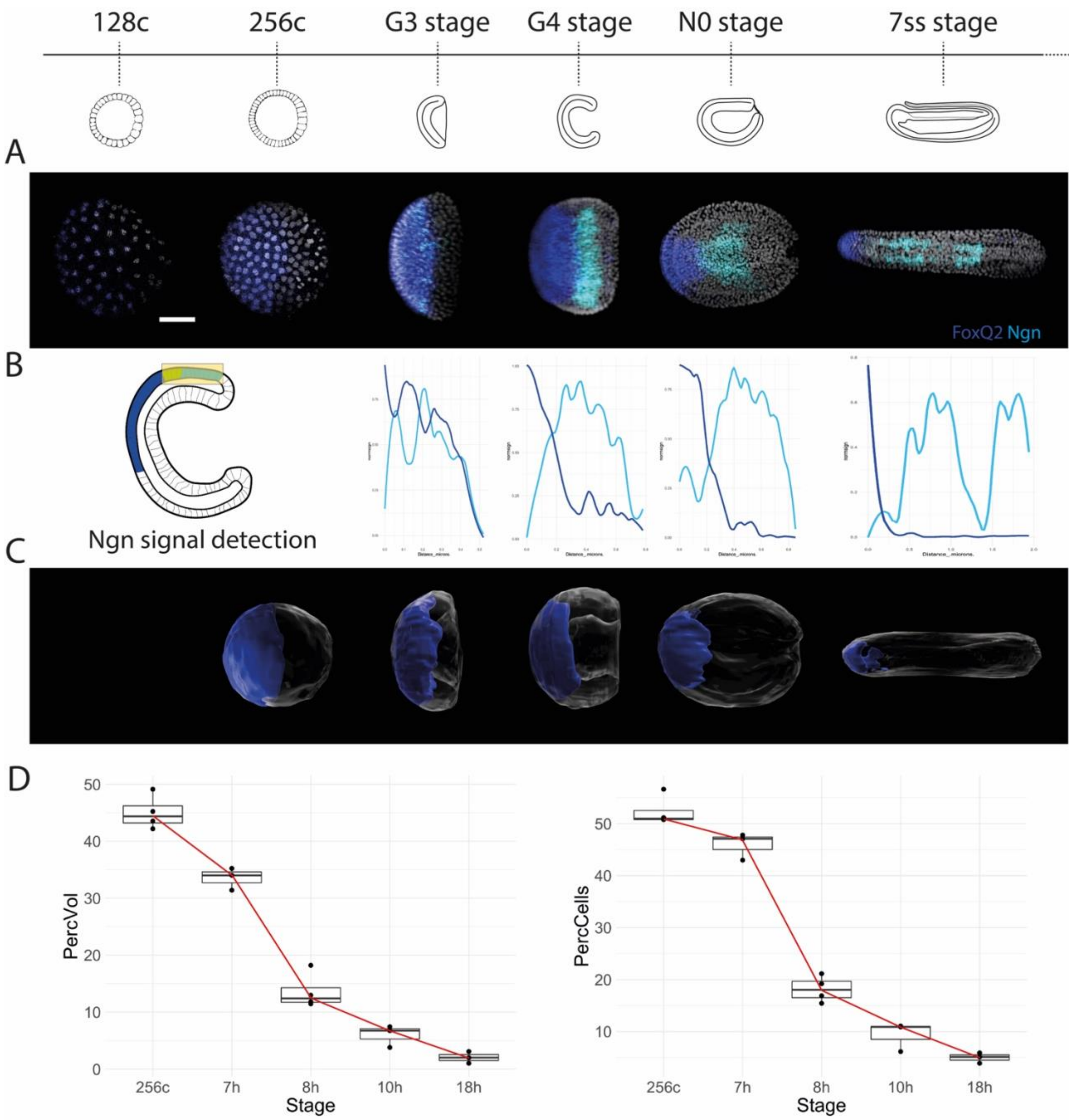


**Supplementary Figure 4. Restriction of FoxQ2 expression during gastrulation.**

**A** Dorsal view of amphioxus embryos from 128 cells to the 7 somites stage showing expression of *FoxQ2* (blue) and *Ngn* (cyan). **B** Distribution of FoxQ2 expression within the *Ngn*-positive neural plate shows restriction of *FoxQ2* outside the neuroectoderm. **C** Manually segmented surfaces of *FoxQ2*-positive ectodermal cells across development. **D** Quantification of *FoxQ2* restriction by counting the percentage of *FoxQ2*-positive cells and of *FoxQ2*-positive volume in embryos at successive stages of development.

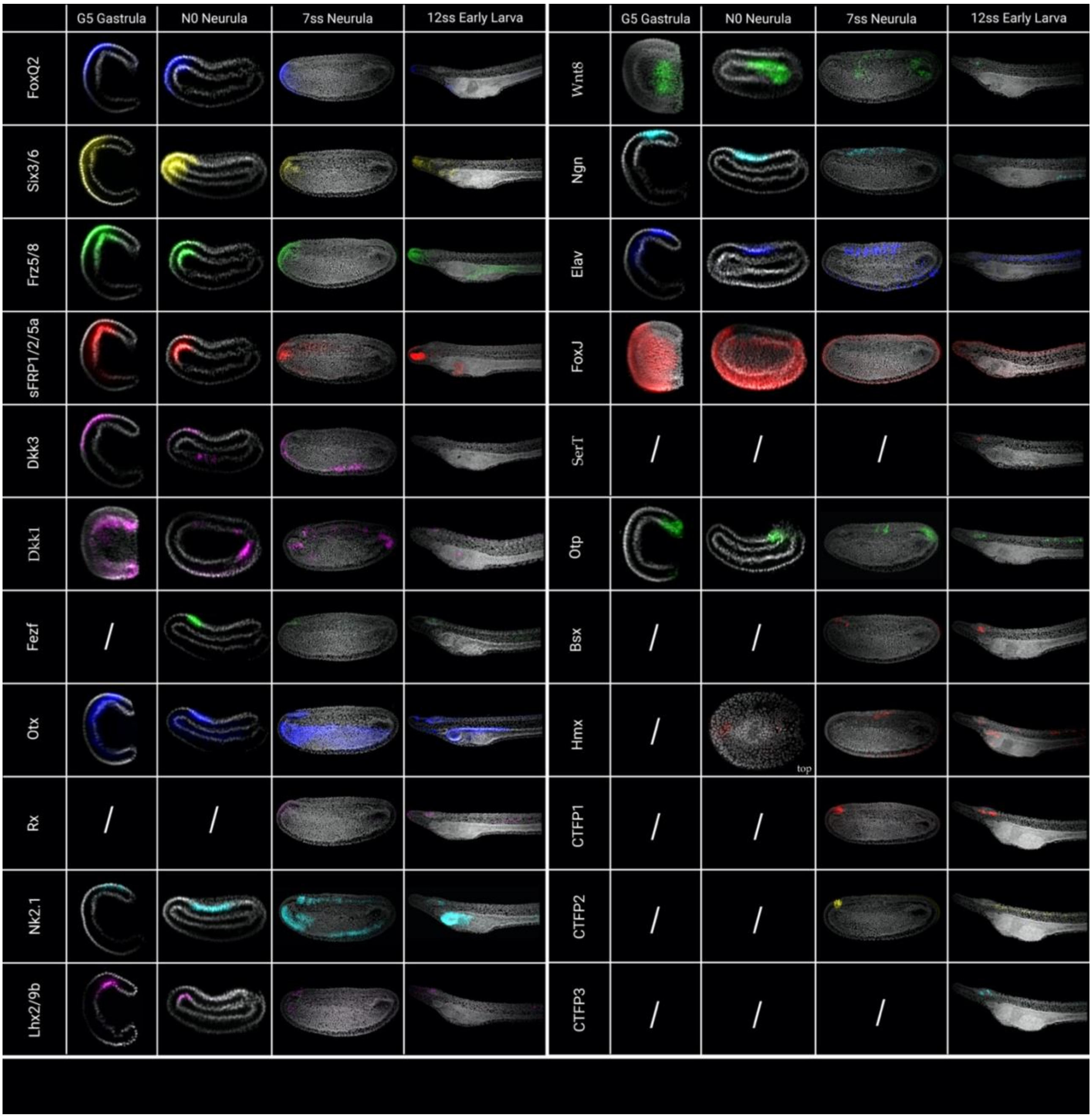


**Supplementary Figure 5. Developmental expression of genes considered in this study visualized through *in situ* HCR in four developmental stages: gastrula (G5), early neurula (N0), mid neurula (7ss) and early larva (12ss).**

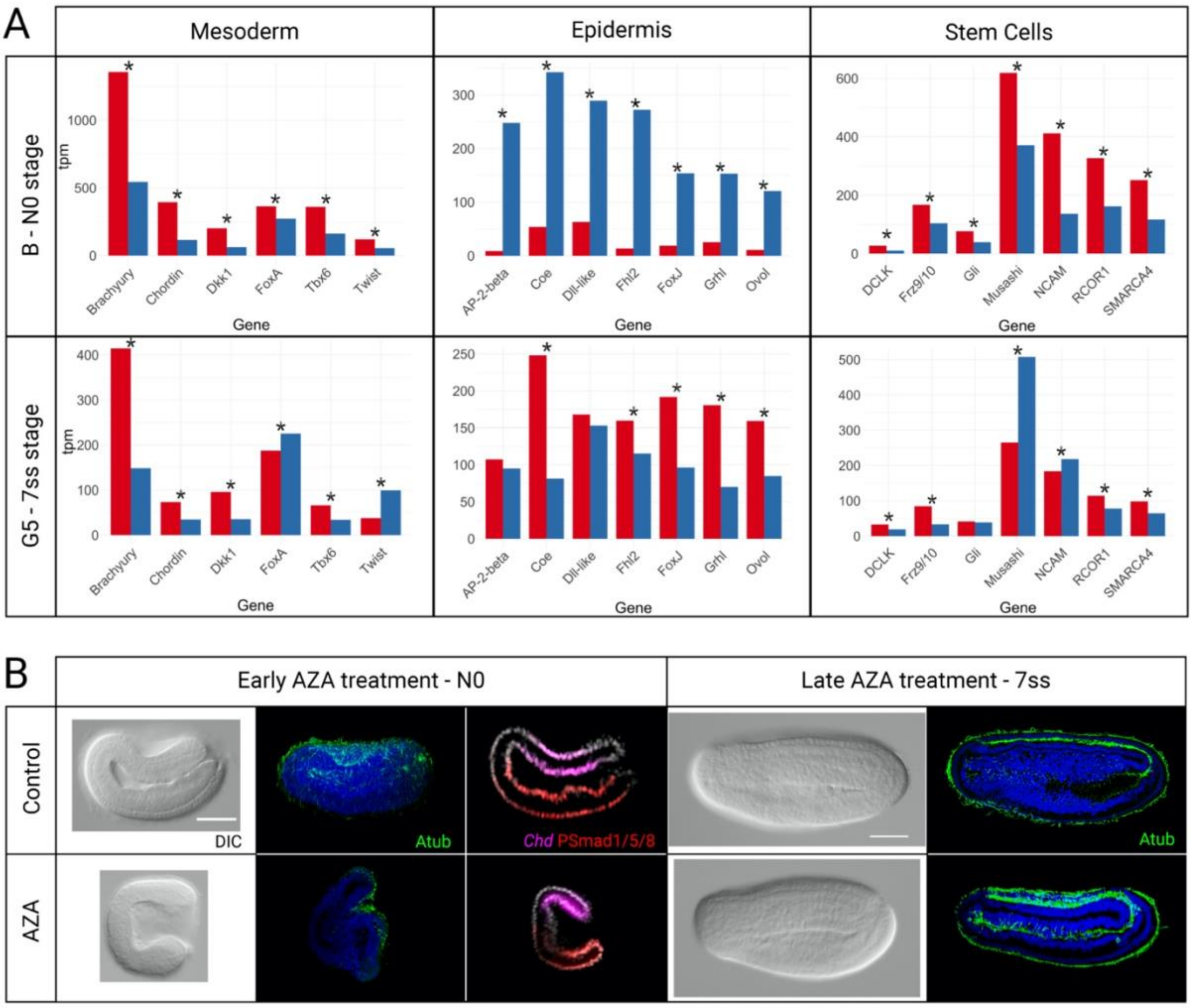


**Supplementary Figure 6. Additional analysis of Azakenpaullone-treated embryos.**

**A** RNAseq analysis of the differential expression in mesodermal, epidermal and stem cell markers in control and Azakenpaullone-treated embryos. **B** Phenotypic characterization in early and late AZA-treated embryos through DIC imaging, Acetylated tubulin (Atub) staining (green), and distribution of dorsal *Chd* expression (magenta) and ventral PSmad2/5/8 (red) immunoreactivity.

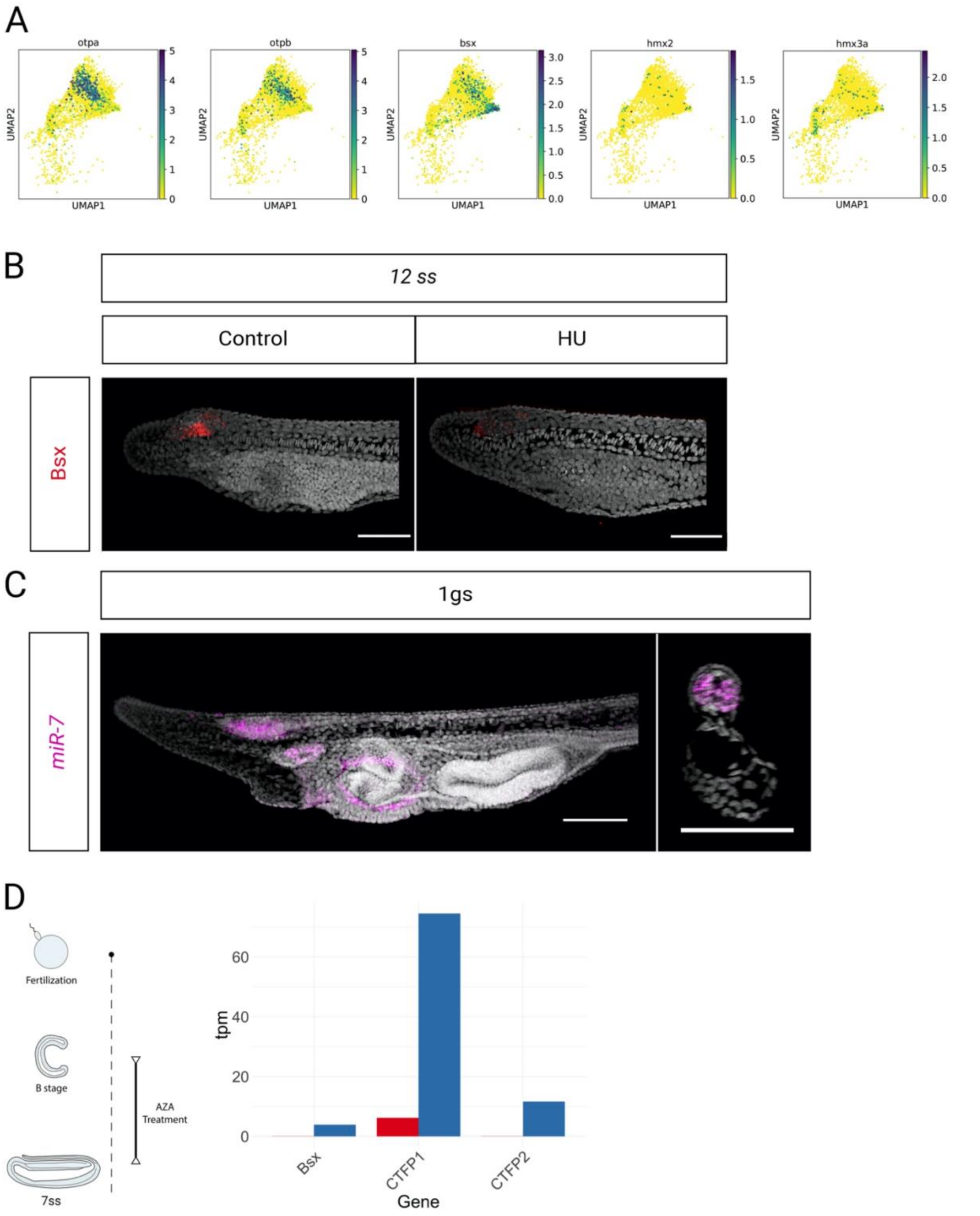


**Supplementary Figure 7. Expression of hypothalamic markers in the amphioxus cerebral vesicle.**
**A** Characterization of hypothalamic markers in zebrafish embryos and larvae. Dataset from Raj et al., 2020. **B** Expression of *Bsx* in the hypothalamus-like region of amphioxus requires neural proliferation: treatment with proliferation blocker Hydroxyurea (HU) from the 7 somite (7ss) stage to the early larva (12ss) leads to loss of the hypothalamic *Bsx* population, while anterior and posterior *Bsx*-positive cells are still visible after treatment. **C** Characterization of the distribution of mature *miR-7* transcripts in

amphioxus larvae (1 gill slit). Transcripts are found within the anterior cerebral vesicle in 1 gill slit larvae. The expression is dorso-lateral while it is absent from the dorsal roof of the brain (see cross section) and from the anteriormost pigment cells. Outside the nervous system, transcripts are found in the preoral pit and around the mouth. Scalebar is 100µm. **D** Loss of aGRN and posteriorization of the brain following late Azakenpaullone treatment led to loss of hypothalamic markers expressed at the 7ss stage (*Bsx*, *CTFP1*, *CTFP2*).

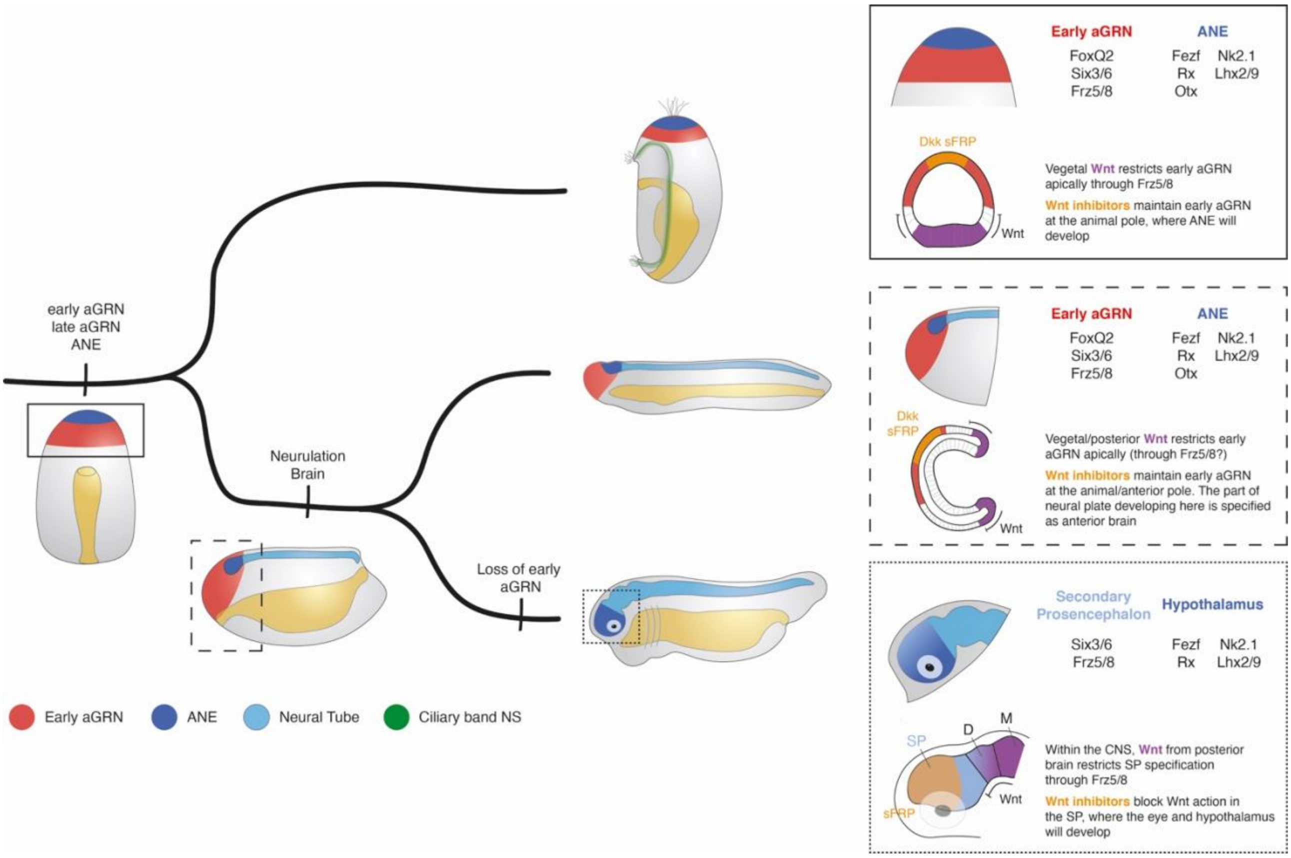


**Supplementary Figure 8. Proposed evolutionary scenario for anterior gene regulatory network (aGRN) and anterior neuroectoderm (ANE) evolution in deuterostomes.**

The last common ancestor of deuterostomes (regardless of his body plan) possessed an aGRN controlling the specification of the ANE on the anterior/apical portion of the body. This aGRN consisted of an early phase, controlled by vegetal Wnt signalling, that defined an apical plate in which Wnt inhibitors were expressed and late phase genes guided the specification of the anterior neuroectoderm (full box). In the ambulacrarian lineage, the ANE develops into the apical organ of the larva. In the chordate lineage, neurogenesis was concentrated on the dorsal side of the embryo, but the aGRN remained active anteriorly, where Wnt inhibitors were expressed, and was still controlled by vegetal Wnt signalling. The aGRN therefore affects only the anterior portion of the neuroectoderm, which acquire a sensory-neurosecretory forebrain identity (dashed box). This condition is still visible in modern cephalochordates. In the vertebrate lineage, the early portion of the aGRN was lost and the network was restricted to the neuroectoderm, where it specified retino-hypothalamic fate (dotted box). Moreover, the main source of Wnt affecting it became the posterior portion of the forebrain.
